# The *9p21.3* Coronary Artery Disease Risk Locus Modulates Vascular Cell-State Transitions via Enhancer-Driven Regulation of *MTAP*

**DOI:** 10.1101/2025.11.18.689066

**Authors:** Timothy N. Audam, Benjamin Schmandt, Yi Huang, Young Beum Cho, Mayank Murali, Rama Kaalia, Saaket Agrawal, Jinping Liu, Hongjie Shi, Yuan Zhou, Dengwei Cao, Lan Jiang, Yun Li, Jichen Wen, Hongqiang Zhao, Alham Saadat, Amelia Hall, Samantha Laber, Sophie Strobel, Siwei Chen, Elizabeth McGonagle, Adam Stefek, Mary Pat Reeve, Alexandra M. Ham, Cassandra M. White, Michelle Melanson, Amit V. Khera, Charles B. Epstein, Mark Chaffin, Patrick T. Ellinor, Minxian Wang, Mark J. Daly, Konrad J. Karczewski, Thouis Jones, Hesam Dashti, Melina Claussnitzer

**Affiliations:** Medical and Population Genetics Program, Cardiovascular Disease Initiative, & Broad Diabetes Initiative, Broad Institute of MIT and Harvard, Cambridge, MA; Center for Genomic Medicine, Massachusetts General Hospital, Harvard Medical School, Boston, MA, USA; The Novo Nordisk Foundation Center for Genomic Mechanism of Disease at Broad Institute, Cambridge, MA, USA of MIT and Harvard; Division of Cardiovascular Medicine, Bringham and Women’s Hospital, Boston, MA, USA; Verve Therapeutics, Boston, MA, USA; Department of Medicine, Harvard Medical School, Boston, MA, USA; Institute for Molecular Medicine Finland (FIMM), Helsinki Institute of Life Science, University of Helsinki, Helsinki, Finland; The University of Chicago, Department of Human Genetics, Chicago, IL, USA; Department of Neuroscience, Scripps Research, La Jolla, CA, USA; China National Center for Bioinformation, Beijing, China; College of Future Technology, Sino-Danish College, University of Chinese Academy of Sciences, Beijing, China; Department of Cardiovascular Surgery, Zhongnan Hospital of Wuhan, Wuhan, China; Hubei Provincial Engineering Research Center of Minimally Invasive Cardiovascular Surgery, Wuhan, China; Wuhan Clinical Center for Minimally Invasive Treatment of Structural Heart Disease, Wuhan, China; Center for the Development of Therapeutics, Broad Institute of MIT and Harvard, Cambridge, MA, USA

## Abstract

The *9p21.3* locus is the strongest genetic association with coronary artery disease (CAD), yet its causal mechanisms remain unresolved. We map the regulatory architecture of *9p21.3* in disease-relevant vascular cells, identifying 12 enhancers within the CAD risk haplotype that respond dynamically to inflammatory and metabolic stress in fibroblasts and smooth muscle cells. These activated states are enriched for CAD heritability, implicating stress-responsive vascular wall cells in disease pathophysiology. Dense CRISPRi tiling integrated with fine-mapping and genomic constraint across >500,000 individuals nominates *MTAP* as the effector gene, with rs1537371 as a likely causal variant. Perturbation and multi-modal analyses show that *MTAP* loss induces pro-fibrotic and angiogenic programs and sensitizes vascular cells to TGF-*β*–driven pathological transitions. Our findings reveal a vascular-specific enhancer network through which noncoding variation at *9p21.3* modulates CAD risk via *MTAP*—a previously unrecognized regulator of vascular remodeling located 269 kb from the risk haplotype.

**Highlights:**

- *9p21.3* CAD risk locus harbors 12 functional enhancers active in vascular wall cell types
- CAD heritability is enriched in stressed vascular fibroblasts and smooth muscle cells
- High-resolution CRISPRi-MAC-seq maps *9p21.3* enhancer-gene and SNP-gene interactions
- *MTAP* identified as a causal effector gene regulated by *9p21.3* fine-mapped and constrained variants
- *MTAP* modulates TGF-β-driven pathological vascular cell transitions

## Main

Despite advances in preventive and interventional strategies, coronary artery disease (CAD) remains the leading global cause of morbidity and mortality ^1,2^. CAD is a complex disease caused by an interplay of both genetic and environmental factors, with narrow sense heritability estimates ranging from 40% to 60% ^3,4^—underscoring a substantial genetic contribution to disease risk.

Large-scale genome-wide association studies (GWAS) have now identified over 240 loci associated with CAD across multiple ancestries ^5–8^. Among these, the chromosome *9p21.3* locus consistently emerges as the most significant genetic risk locus for CAD, particularly in non-African populations ^9–12^. Beyond CAD, *9p21.3* has been associated with other complex diseases such as type 2 diabetes ^13^, abdominal aortic and intracranial aneurysm ^14^, and cancer ^15^. Despite more than a decade of research, the mechanistic link between *9p21.3* genetic variation and disease-causing mechanisms remains elusive.

Recent studies have begun to explore the regulatory potential of the *9p21.3* disease risk locus ^16–19^. Harismendy et al. (2011) identified candidate enhancer elements at *9p21.3* with allele-specific activity in vitro ^20^, while Lo Sardo et al. (2018) modeled the locus using haplotype editing in iPSC-derived smooth muscle cells and reported regulation of risk-dependent gene networks driven by Antisense Noncoding RNA in the INK4 Locus (ANRIL) perturbation ^21^. More recently, Salido et al. (2025) showed that the risk haplotype in iPSC-derived vascular smooth muscle cells activated an osteochondrogenic program ^22^. In mice, deletion of the orthologous ∼70-kb region perturbs *Cdkn2a/b* expression but fails to elicit atherosclerotic phenotypes, raising concerns about mechanistic conservation across species ^23^. Collectively, these studies highlight regulatory activity at the locus and support a role in smooth muscle cells. However, prior studies have neither mapped and explained the effect of CAD disease-driving SNPs at the *9p21.3* locus, nor comprehensively mapped variant-to-enhancer-to-gene connections across CAD related vascular cell types and their disease relevant stimulated cell states. Our study addresses this by systematic epigenomic profiling, dense CRISPRi-based tiling, and fine-mapping across multiple vascular cell types, generating the first comprehensive map of functional variant-to-gene connections at *9p21.3*.

The *9p21.3* locus is a highly complex locus spanning a 50-kb interval embedded within the long non-coding RNA CDKN2B-AS1, encompassing ∼ 59 SNPs in high linkage disequilibrium (LD). These variants are located about 100-kb upstream of the cell cycle repressor coding genes *CDKN2A/CDKN2B* ^23,24^. Multiple studies have proposed that the locus may interact with other risk factors—including aging ^16^, diet ^25^, hypercholesterolemia ^26^, and type 2 diabetes ^27^— while others suggest its effects are largely independent of traditional CAD risk pathways ^28^. Still, the identity of the effector transcript(s), the relevant cell type(s), and the disease-context–dependent mechanisms through which *9p21.3* confers CAD risk remain unclear ^22^.

To dissect the regulatory logic of the *9p21.3* locus, we constructed a multi-layered functional genomic framework that integrates reference epigenomes of vascular wall cell types with CRISPR interference tiling, statistical fine-mapping, 3D chromatin conformation, and deep image-based phenotyping. This integrative strategy enabled us to map an atlas of disease-associated variants to functional enhancers in disease-relevant cell types, connect enhancers to target genes, and nominate a causal effector gene—*MTAP*—mediating vascular remodeling and CAD susceptibility.

## Results

### Phenome-Wide Associations Study Links *9p21.3* to Vascular Wall-Related Disease Processes

Although CAD primarily affects the coronary vasculature — including endothelial and smooth muscle cells — its clinical consequences involve diverse cardiac pathophysiologies such as cardiomyocyte dysfunction, arrhythmia, mitochondrial impairment, and inflammation. To identify the primary disease processes linked to *9p21.3* variation ^29^, we queried the results of a phenome-wide association study (PheWAS) in FinnGen data freeze 13 (DF13), comprising over 500,000 individuals and 2,400 clinical phenotypes ^30^. The analysis revealed a strikingly specific pattern of association between the *9p21.3* locus and vascular wall-related disease traits (**Figure 1A**; **Table S1**). We observed the strongest associations with atherosclerotic vascular conditions and interventional cardiovascular procedures, including coronary atherosclerosis (β=0.18; P-value=1.48×10^-159^), coronary revascularization (ANGIO or CABG) (β=0.14; P-value=5.64×10^-116^), coronary artery bypass grafting (β=0.35; P-value=9.12×10^-154^), and percutaneous coronary intervention (angioplasty) (β=0.13; P-value= 3.58×10^-107^) — all procedures that directly address atherosclerotic vascular pathology. Additional top PheWAS signals included major CAD endpoints, including ischemic heart disease (β=0.15; P-value=3.87×10^-138^), myocardial infarction (β= 0.19; P-value=1.39×10^-106^), both stable (β=0.17; P-value=3.58×10^-107^) and unstable (β=0.19; P-value=2.60×10^-62^) angina pectoris, and major coronary heart disease events (β=0.13; P-value=1.04×10^-114^). Notably, associations extended to systemic vasculopathies, including peripheral atherosclerosis (β=0.12, P-value=1.76×10⁻^35^), small artery and capillary disease (β=0.13, P-value=2.02×10⁻^35^), abdominal aortic aneurysm (β=0.19, P-value=1.35×10⁻^19^), and cerebrovascular phenotypes (e.g., stroke, subarachnoid hemorrhage). Retinal vascular diseases — such as glaucoma and macular degeneration — were also enriched, supporting a model in which independent genetic signals at *9p21.3* influence shared vascular wall processes rather than vessel bed-specific mechanisms (**Table S1**).

**Figure 1.**
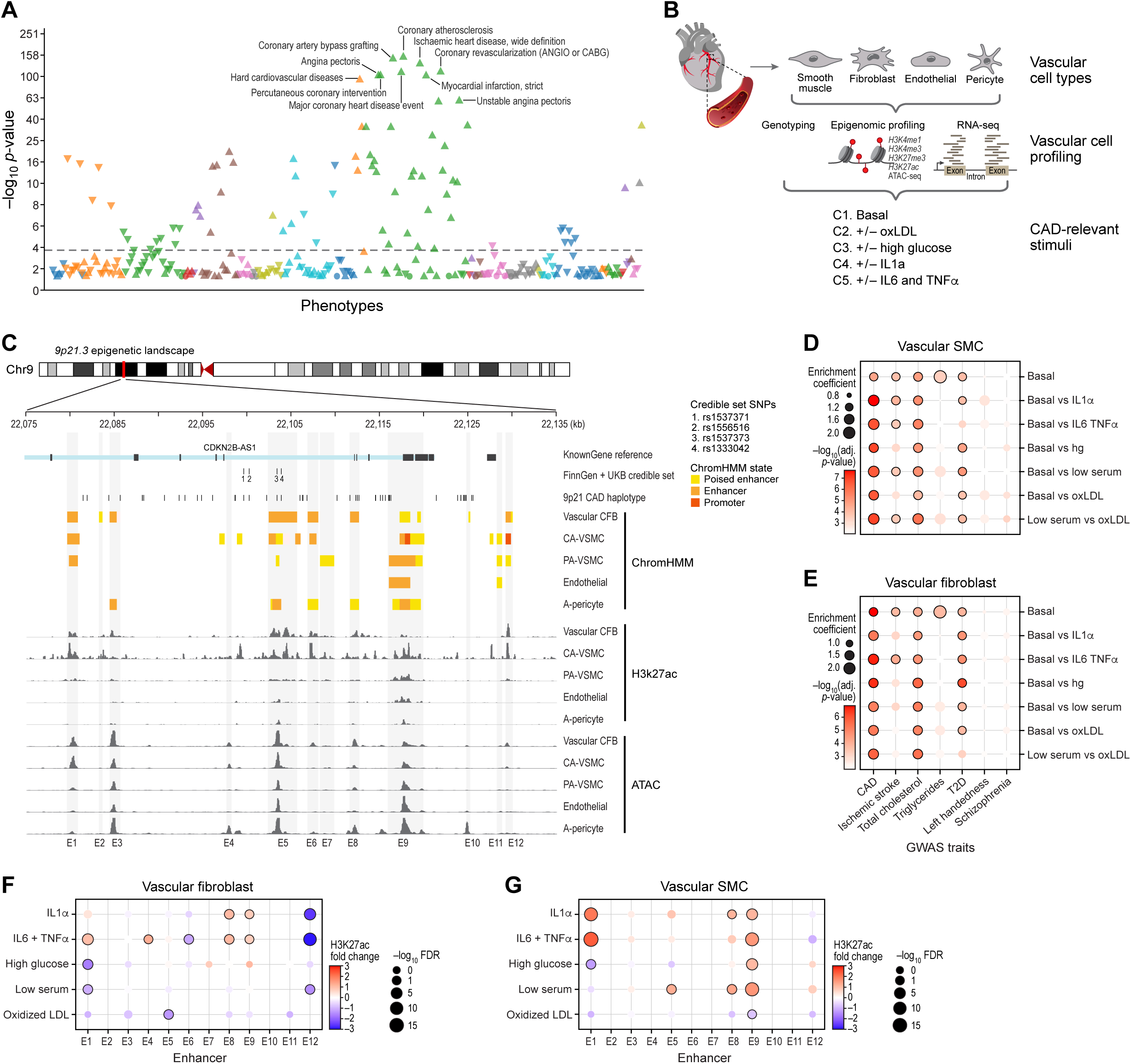
The *9p21.3* coronary artery disease (CAD) locus maps to an enhancer dense region in CAD-relevant cell types. (A) Manhattan plot showing Phenome-wide association (PheWAS) of *9p21.3* locus (tagged by rs4977574) to multiple disease traits in FinnGen data freeze 13 (DF13). Each triangle represents a phenotype association at *9p21.3* locus and different triangle colors represent various classification of phenotypes. Direction of the triangle indicates increased directionality of association. Increased risk is indicated by an upward triangle and reduced risk is indicated by a downward facing triangle. (B) Schematic depicting an extensive vascular cell wall profiling at baseline and in response to CAD-relevant stimulatory condition. (C) Chromatin state and epigenetic landscape of *9p21.3* CAD haplotype spanning ∼ 50-kb DNA sequence within the 3’ region of long non-coding RNA *CDKN2B-AS1* (*ANRIL*). Also depicted are all 50 CAD associated SNPs (r^2^ ≥ 0.8), tagged by rs4977573 and credible set variants from FinnGen and UKBB finemapping efforts. (D-E) Dot plots showing enrichment for CAD heritability at baseline and in response to CAD-relevant stimulatory conditions, including Inflammatory (IL-1α and IL-6 & TNF-α) vascular fibroblast and smooth muscle cells. Solid dark lines around each dot represent significance at a threshold of -log_10_(adjusted P-value) ≥ 3.35. (F-G) Dot plots showing the impact of CAD-relevant stimulatory conditions on enhancer activities (proxy of H3K27ac) at *9p21.3* locus in vascular fibroblast and smooth muscle cells. Fold changes are indicated as Red (increased fold change) or Blue (Reduced fold change) and solid dark lines around each dot represent significance at a threshold of -log_10_(FDR) ≥ 1.3.

In addition to vascular phenotypes, the locus was modestly associated with upstream cardiometabolic risk factors: disorders of lipoprotein metabolism (β=0.06, P-value=3.0×10⁻⁴), hypercholesterolemia (β=0.05, P-value = 7.8×10⁻^16^), type 2 diabetes (β=0.03, P-value=1.06×10⁻⁷), and hypertensive heart disease (β=0.04, P-value= 0.0012). Importantly, no associations were observed with primary myocardial disorders such as dilated or hypertrophic cardiomyopathy. We also observed nominal signals for heart failure (β=0.04, P-value= 3.92×10⁻^6^) and atrial fibrillation (β=0.08, P-value=2.46×10⁻^17^) – which likely reflect downstream effects of vascular disease.

Although many of these aforementioned traits show genetically distinct association from CAD association, these findings indicating specificity of vascular associations point to vascular wall–centric mechanisms as the primary driver of *9p21.3*-mediated CAD risk.

### Mapping The *9p21.3* Locus To Regulatory Elements In Disease-Relevant Vascular-Wall Cell Type

The *9p21.3* locus spans ∼50-kb DNA sequence within the 3’ region of a long non-coding RNA *CDKNB-AS1* ^24^ and includes 50 non-coding variants in strong LD (r^2^ ≥ 0.8), tagged by rs4977574 (**Table S2**). Given the vascular specificity of *9p21.3*-associated traits, we sought to map these variants to functional regulatory elements in disease-relevant vascular wall cells.

First, we examined chromatin state maps across 111 cell types and tissues from the Roadmap Epigenome ^31^ and ENCODE consortia ^32^ (**Figure S1A**). These publicly available chromatin state maps appeared to lack adequate vascular cell representation beyond Human Umbilical Vein Endothelial Cells (HUVECs) and aortic smooth muscle cells (SMCs) ^31^. To fill this gap, we generated universal chromatin state maps across five primary human vascular wall cell types: vascular fibroblasts (atrial-derived), vascular smooth muscle cells (pulmonary and coronary artery-derived), endothelial cells (coronary artery-derived), and pericytes (adipose-derived) (**Figure 1B**). We employed Mint-ChIP, a barcoded and multiplexed chromatin profiling technology that enables quantitative histone mark mapping from low-input samples via indexed T7 amplification and pool-and-split library generation ^33^. This platform allowed high-throughput, quantitative profiling of key histone modifications marking enhancers (histone 3 lysine 4 monomethylation, H3K4me1 and histone 3 lysine 27 acetylation, H3K27ac) ^34^; promoters (histone 3 lysine 4 trimethylation, H3K4me3) ^35^; and polycomb-repressed chromatin (histone 3 lysine 27 trimethylation, H3K27me3) ^36^, alongside chromatin accessibility (ATAC-seq). Transcriptomes (RNA-seq) and genotyping were also collected (**Figure 1B**). To segment reference epigenomes across vascular wall cell types, we used a multivariate hidden Markov model, ChromHMM, which explicitly models the combinatorial presence or absence of each of the profiled chromatin marks ^37^. Using ChromHMM, we inferred 18 chromatin states per vascular wall cell types (**Figure S1B-1H**), yielding annotated epigenomes of cells in the vascular wall. This analysis identified a total of 774,011 enhancers, including 29,397 novel enhancers not previously annotated in any of the 111 Roadmap reference epigenomes (**Figure S1I**, see Methods). These data reveal a rich landscape of genome-wide vascular wall-specific gene regulation, enabling direct mapping of GWAS loci in relevant cellular contexts.

Intersecting these enhancer maps along with ATAC-seq and H3K27ac ChIP-seq data with the *9p21.3* CAD haplotype (**Table S2**) ^38^ revealed a dense cluster of 12 enhancers (E1-12), suggesting a role of the *9p21.3* disease locus in vascular wall regulatory activities (**Figure 1C**). Many of the 50 *9p21.3* CAD haplotype variants overlap or are flanking the annotated enhancers, with notable concentration flanking enhancer E4 and within enhancer E5 near a series of fine-mapped credible set variants from FinnGenn and UKBB (**Table 1**). Annotated enhancer E5 showed particularly strong H3K27-acetylation activity in both vascular fibroblasts and vascular smooth muscle cells (**Figure 1C**) compared to other vascular cell types. While this enhancer-dense region exhibited some characteristics of a super-enhancer ^39^, formal analyses using the Rank Ordering of Super Enhancer (ROSE) algorithm ^40,41^ with H3K27ac signals as inputs found no evidence of a unified super-enhancer structure in vascular cells (**Figure S2A-2D**). Rather, the *9p21.3* CAD risk locus forms a modular enhancer hub in vascular wall cells, potentially capable of integrating environmental cues and genetic signals potentially through vascular cell-specific regulatory programs.

**Table 1.** Fine-mapped variants associated with a compbined trait for coronary revascularization, including coronary artery bypass grafting (CABG) and percutanutous coronary intervention (ANGIO) using data from large-scale population efforts from FinnGen and UKBB.

| chr | pos | ref | alt | var | rsID | FG_beta | FG_se | FG_p | FG_AF | UK_beta | UK_se | UK_p | UK_AF | meta_beta | meta_se | meta-logp |
| --- | --- | --- | --- | --- | --- | --- | --- | --- | --- | --- | --- | --- | --- | --- | --- | --- |
| 9 | 22088091 | A | T | 9:22088091:A:T | rs10738606 | 2.86E-01 | 9.76E-03 | 1.17E-188 | 0.4107 | 2.54E-01 | 1.08E-02 | 1.12E-121 | 0.4851 | 2.72E-01 | 7.25E-03 | 306 |
| 9 | 22088095 | A | G | 9:22088095:A:G | rs10738607 | 2.86E-01 | 9.76E-03 | 1.10E-188 | 0.4107 | 2.54E-01 | 1.08E-02 | 1.05E-121 | 0.4851 | 2.72E-01 | 7.25E-03 | 307 |
| 9 | 22088261 | C | T | 9:22088261:C:T | rs10757272 | 2.87E-01 | 9.77E-03 | 3.00E-189 | 0.41 | 2.54E-01 | 1.08E-02 | 3.75E-122 | 0.4842 | 2.72E-01 | 7.25E-03 | 308 |
| 9 | 22090522 | G | A | 9:22090522:G:A | rs9644859 | 2.88E-01 | 9.75E-03 | 6.71E-192 | 0.4169 | 2.60E-01 | 1.08E-02 | 8.89E-127 | 0.4939 | 2.75E-01 | 7.25E-03 | 315 |
| 9 | 22090604 | C | T | 9:22090604:C:T | rs9644860 | 2.88E-01 | 9.75E-03 | 7.52E-192 | 0.4169 | 2.60E-01 | 1.08E-02 | 1.02E-126 | 0.4939 | 2.75E-01 | 7.25E-03 | 315 |
| 9 | 22090936 | C | T | 9:22090936:C:T | rs9644861 | 2.88E-01 | 9.75E-03 | 7.57E-192 | 0.4169 | 2.61E-01 | 1.08E-02 | 2.86E-128 | 0.4942 | 2.76E-01 | 7.25E-03 | 317 |
| 9 | 22090937 | T | G | 9:22090937:T:G | rs9644862 | 2.79E-01 | 9.71E-03 | 1.39E-181 | 0.4336 | 2.61E-01 | 1.08E-02 | 2.80E-128 | 0.4942 | 2.71E-01 | 7.23E-03 | 307 |
| 9 | 22091070 | C | T | 9:22091070:C:T | rs10811653 | 2.88E-01 | 9.75E-03 | 6.70E-192 | 0.4169 | 2.60E-01 | 1.08E-02 | 8.65E-127 | 0.4939 | 2.75E-01 | 7.25E-03 | 315 |
| 9 | 22099569 | C | A | 9:22099569:C:A | rs1537371 | 2.90E-01 | 9.74E-03 | **2.45E-195** | 0.4201 | 2.62E-01 | 1.08E-02 | **8.41E-129** | 0.4958 | 2.78E-01 | 7.24E-03 | **320** |
| 9 | 22100177 | G | C | 9:22100177:G:C | rs1556516 | 2.91E-01 | 9.74E-03 | **1.14E-195** | 0.4201 | 2.62E-01 | 1.08E-02 | **7.83E-129** | 0.4959 | 2.78E-01 | 7.24E-03 | **321** |
| 9 | 22102166 | C | T | 9:22102166:C:T | rs7859727 | 2.88E-01 | 9.75E-03 | 3.16E-192 | 0.4132 | 2.59E-01 | 1.08E-02 | 4.41E-126 | 0.4872 | 2.75E-01 | 7.25E-03 | 315 |
| 9 | 22103342 | T | G | 9:22103342:T:G | rs1537373 | 2.90E-01 | 9.74E-03 | 1.15E-194 | 0.4195 | 2.62E-01 | 1.08E-02 | **5.71E-129** | 0.4952 | 2.77E-01 | 7.24E-03 | **320** |
| 9 | 22103814 | A | G | 9:22103814:A:G | rs1333042 | 2.91E-01 | 9.74E-03 | **1.26E-195** | 0.42 | 2.63E-01 | 1.08E-02 | **1.01E-129** | 0.4962 | 2.78E-01 | 7.24E-03 | **322** |
| 9 | 22123767 | A | C | 9:22123767:A:C | rs10738610 | 2.85E-01 | 9.77E-03 | 8.07E-188 | 0.4184 | 2.55E-01 | 1.08E-02 | 1.42E-122 | 0.4932 | 2.72E-01 | 7.26E-03 | 307 |
| 9 | 22124124 | T | A | 9:22124124:T:A | rs1333046 | 2.84E-01 | 9.75E-03 | 6.92E-187 | 0.4199 | 2.55E-01 | 1.08E-02 | 1.00E-122 | 0.496 | 2.71E-01 | 7.25E-03 | 306 |
| 9 | 22124141 | A | T | 9:22124141:A:T | rs7857118 | 2.86E-01 | 9.74E-03 | 1.10E-189 | 0.426 | 2.60E-01 | 1.09E-02 | 7.10E-127 | 0.5052 | 2.74E-01 | 7.25E-03 | 313 |
| 9 | 22124451 | A | G | 9:22124451:A:G | rs10757277 | 2.86E-01 | 9.75E-03 | 3.59E-189 | 0.4186 | 2.52E-01 | 1.08E-02 | 4.02E-120 | 0.4799 | 2.71E-01 | 7.25E-03 | 305 |
| 9 | 22124478 | A | G | 9:22124478:A:G | rs10757278 | 2.86E-01 | 9.75E-03 | 3.24E-189 | 0.4184 | 2.52E-01 | 1.08E-02 | 1.36E-119 | 0.4801 | 2.71E-01 | 7.25E-03 | 305 |
| 9 | 22124505 | A | T | 9:22124505:A:T | rs1333047 | 2.88E-01 | 9.74E-03 | 4.59E-192 | 0.4245 | 2.56E-01 | 1.08E-02 | 8.05E-124 | 0.4891 | 2.74E-01 | 7.24E-03 | 312 |
| 9 | 22124631 | A | G | 9:22124631:A:G | rs10757279 | 2.86E-01 | 9.75E-03 | 3.61E-189 | 0.4186 | 2.52E-01 | 1.08E-02 | 3.20E-120 | 0.48 | 2.71E-01 | 7.25E-03 | 305 |
| 9 | 22124745 | C | G | 9:22124745:C:G | rs4977575 | 2.84E-01 | 9.74E-03 | 2.52E-187 | 0.4307 | 2.56E-01 | 1.08E-02 | 1.05E-123 | 0.4894 | 2.72E-01 | 7.24E-03 | 307 |
| 9 | 22125348 | A | C | 9:22125348:A:C | rs1333048 | 2.84E-01 | 9.75E-03 | 8.24E-187 | 0.4197 | 2.55E-01 | 1.08E-02 | 1.21E-122 | 0.4962 | 2.71E-01 | 7.25E-03 | 306 |

### CAD Heritability and 9p21.3 Regulatory Activity Is Enriched in Stimulated Vascular Fibroblasts and Vascular Smooth Muscle Cells

Connecting genetic variation to disease physiology is challenging since biological networks are complex, and cellular programs are dynamic and stochastic. Prior studies have shown that many non-coding variants that are not associated with gene expression levels at steady state are found to affect dynamic programs of gene expression in specific disease-relevant contexts ^42,43^, and *9p21.3* CAD risk was found to be largely increased when patients present with poor glycemic control ^44^.

To identify the vascular cell types and states in which CAD risk variation may become functionally active, we exposed our primary human vascular cells to a range of CAD-relevant stimuli. These included inflammatory cytokines (IL-1α, IL-6 andTNF-α), oxidized LDL (oxLDL) that model the inflammatory environment of atherosclerotic plaques, and metabolic stressors (high glucose and low serum) that reflect diabetic vascular dysfunction. We then profiled both RNA-sequencing along with epigenomic marks enriched at enhancers including H3K27ac and chromatin accessibility (ATAC-seq) across all perturbations (**Figure 1B**). To prioritize which vascular wall cell types and states are most relevant for mediating CAD risk, we first partitioned the heritability of CAD and related traits using stratified LD score regression ^45^ using GWAS summary statistics and transcriptional profiles of vascular cells. We observed significant enrichment for CAD heritability particularly in vascular fibroblasts and vascular smooth muscle cells under inflammatory and metabolic stimulations (**Figure 1D–1E**; **Table S3**). For instance, fibroblasts treated with IL-6+TNF-α or high glucose showed the highest enrichment coefficients (1.75, P-adj=2.1×10⁻7 and 1.50, P-adj=8×10⁻^7^, respectively). Similarly, CAD heritability was enriched in smooth muscle cells stimulated with IL-1α (enrichment coefficient= 1.8, P-adj= 2.3×10^-8^), IL-6+TNF-α (enrichment coefficient=1.8, P-adj=2.3×10^-7^), and high glucose (enrichment coefficient= 1.5, P-adj=1.1×10^-7^). These effects were reduced in unstimulated (basal) conditions (enrichment coefficient= 1.3, P-adj= 2.2×10^-5^) (**Table S3**), highlighting the importance of modeling disease-relevant environments. We did not observe any notable enrichment of heritability of unrelated traits such as lefthandedness and schizophrenia, confirming the specificity of our findings (**Figure 1D–1E; Figure S2E–G; Table S3**).

To determine whether these CAD-relevant stimulatory conditions alter enhancer activity at *9p21.3*, we quantified changes in H3K27ac across the ∼50-kb haplotype region in each stimulus condition. Multiple enhancers showed increased acetylation in fibroblasts and smooth muscle cells exposed to inflammatory cytokines or metabolic stress (FDR<0.05; **Figure 1F–G**; **Table S4**). Endothelial cells and pericytes showed modest changes across all H3K27ac peaks, further supporting fibroblasts and smooth muscle cells as key mediators of genetic risk at this locus (**Figure S2H-S2I**, **Table S4).**

Together, these data demonstrate that the *9p21.3* enhancer cluster functions as a context-sensitive regulatory hub, responsive to pro-atherogenic signals particularly in vascular fibroblasts and smooth muscle cells. We therefore prioritized these two cell types for functional dissection of enhancer activity and gene regulation at the *9p21.3* locus.

### Functional Enhancer Dissection of the *9p21.3* Locus in Vascular Fibroblasts and Smooth Muscle Cells

To systematically map functional enhancers and their target gene(s) at the non-coding *9p21.3* locus, we implemented a scalable CRISPR interference (CRISPRi) tiling screen coupled to a multiplexed read-out using MAC-seq ^46^ (CRISPRi-MAC-seq), a cost-efficient technology that allows for simultaneous measurement of gene expression changes across thousands of perturbations while maintaining single-cell-level sensitivity. This multiplexed approach dramatically increased experimental throughput, and therefore enabled unbiased interrogation of enhancer-to-gene (E2G) connections across the 50-kb noncoding risk haplotype in disease-relevant vascular cell types.

We first generated stable dCas9-KRAB-expressing human vascular smooth muscle cells (SMCs) and human vascular fibroblasts (CFBs) ^47^. To validate the fidelity of our engineered cell lines, we assessed mRNA and protein expression levels of dCas9 in our dCas9-KRAB-CFBs and dCas9-KRAB-SMCs, confirming robust dCas9 mRNA and protein level expression compared to parental controls using qPCR and western blot (**Figure S3A-C**), and validated a robust repression activity (>75% efficiency) using flow cytometry-based assays (**Figure S3D-S3E**, see methods).

To delineate the topologically constrained regulatory architecture of the *9p21.3* disease risk locus, we first analyzed 3D genome conformation data from micro-C data ^48^, which revealed the physical boundaries of the locus: a 1 Mb topologically associating domain (TAD) with two sub-TADs (**Figure 2A**). The proximal sub-TAD, containing the *9p21.3* risk haplotype, encompasses several protein-coding genes including the cell cycle regulators *CDKN2B and CDKN2A,* and the long non-coding RNA (*CDKN2B-AS1*), and more genes located further away from the *9p21.3* risk variants including the metabolic enzyme *MTAP,* and the transcription factor *DMRTA1.* The distal sub-TAD contains the micro-RNA gene *MIR31HG*, and multiple interferon alpha genes (*IFNA1*, *IFNA2*, and *IFNA17*) (**Table S5**).

**Figure 2.**
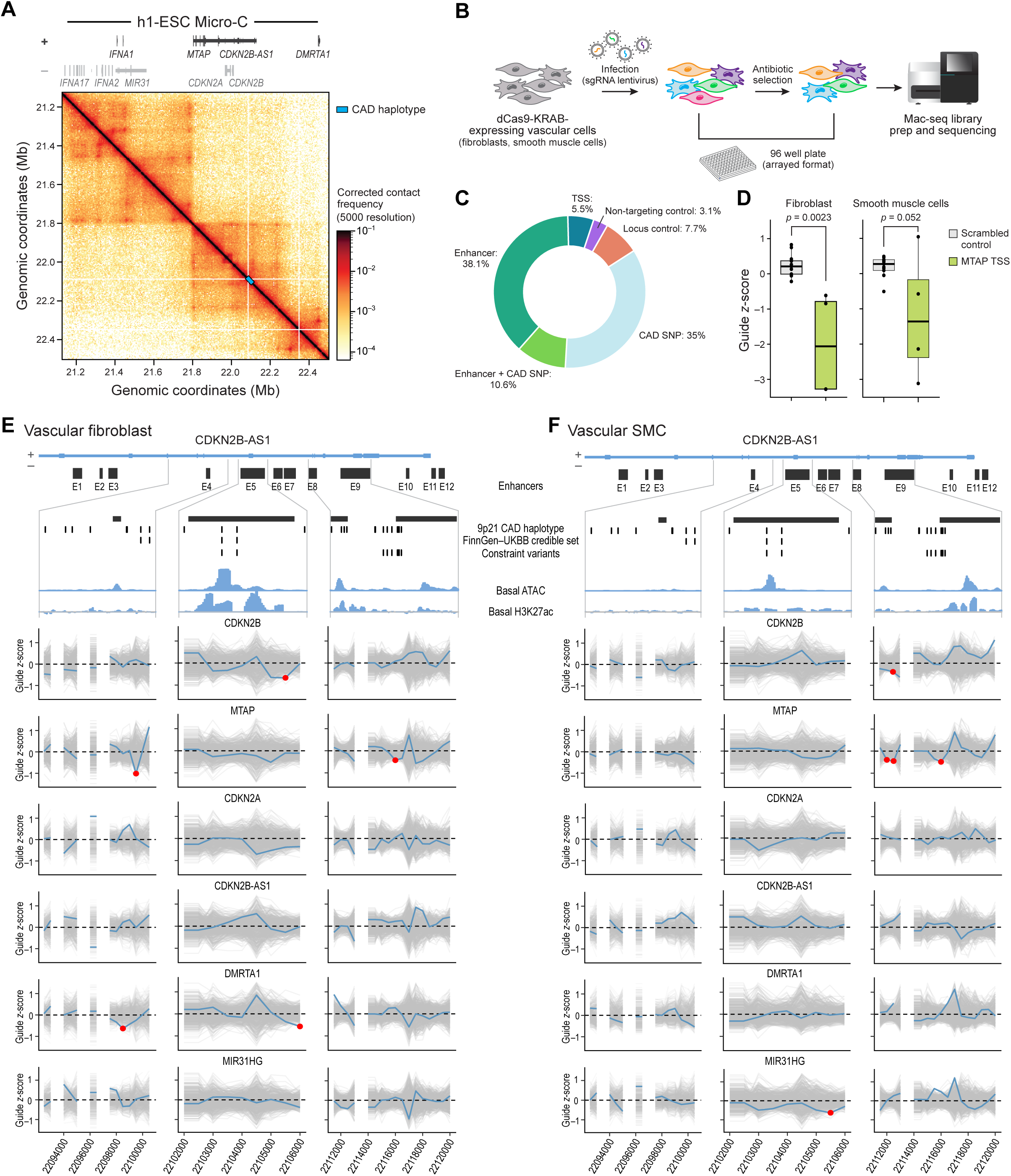
The *9p21.3* locus contains multiple functional vascular cell enhancers regulating cis-expressing genes. (A) Contact matrix (at 5000 bp resolution) from Micro-C^1^ analysis of the H1 human embryonic stem cell (h1-ESC) spanning 1 Mb around *9p21.3* CAD haplotype (indicated as a blue rectangle). (B) Schematic showing experimental workflow to identify all functional non-coding *9p21.3* elements by CRISPR interference (CRISPRi) paired with Multiplex Analysis of Cells (MAC-seq) sequencing readout collectively referred to as CRISPRi-MAC-seq. (C) Donut plot showing the proportion of control, TSS-targeting, enhancer-targeting, and SNP-targeting guides (n=417 total guides) in our densely-tiled *9p21.3* library. (C) Box plots showing TSS repression efficiency via assessing normalized *MTAP* expression in vascular fibroblasts and smooth muscle cells treated with lentivirus containing transcriptional start site (TSS) guides for MTAP (n=4) compared to non-targeting control (n=13). Significance was determined using a two-tailed t-test, with a significance threshold of P-value<0.05. (E-F) Plot shows a sliding window approach that aggregates sgRNA transcriptional effects across a subset of the *9p21.3* annotated enhancers, including E4, E5, E8 and E9 in vascular fibroblast and smooth muscle cells. To assign significance to observed data relative to the permutation background, a permutation z-score was calculated. A significance level of z-score<-1.96 was used to identify significantly repressed regions – illustrated as red points on the sliding window plots.

To enable systematic fine-mapping of the *9p21.3* CAD disease locus, we designed a dense CRISPRi sgRNA library that tiles the 50-kb non-coding haplotype region. The library comprised 417 guides in total (**Table S6**) with 38.1% of guides targeting our annotated vascular enhancers, 35% of guides targeting *9p21.3* CAD associated SNPs, 10.6% of guides targeting enhancers overlapping CAD SNPs, 5.5% of guides targeting transcriptional start sites (TSSs) of *9p21.3* candidate target genes, 3.1% non-targeting control guides, and 7.7% internal locus controls – devoid of any epigenetic marks or chromatin state annotation (**Figure 2C**; **Table S7**).

Following lentiviral delivery of infection using our densely tiled *9p21.3* packaged lentivirus library (**Figure 2B**), CRIPSRi-MAC-seq enabled comprehensive transcriptome profiling to quantify gene expression changes across perturbations including both proximal gene effects and long-range regulatory interactions across sub-TADs. We first validated the sensitivity of our approach by confirming efficient knockdown of candidate target genes in both dCas9-KRAB-CFBs (**Figure 2D**; **Figure S4A–4D**) and dCas9-KRAB-SMCs (**Figure 2D**; **Figure S4E–4H**), with guide-dependent knockdown detection and some degree of inter-replicate correlations of QC parameters – similar to previously reported large CRISPRi study ^49^ (**Figure S10A-10D**).

Our CRISPRi-MAC-Seq strategy revealed extensive enhancer-to-gene (E2G) connections with strong cell type specificity (**Figure S3F-3G**). We calculated a normalized effect z-score by dividing the mean effect on transcription for each gene by the standard deviation of sgRNA effects from the non-targeting control set as previously described ^50^. In vascular fibroblasts, we identified robust regulatory effects of E3, and E5–E12 on *CDKN2B*, and of E5, E7–E9 on *MTAP* (**Figure S3F**). Enhancers E3–E4 also regulated *DMRTA1*, enhancer E3, E7-E8 regulated *MIR31HG*, while *CDKN2B-AS1* showed minimal enhancer connectivity. In vascular smooth muscle cells, *CDKN2A* and *CDKN2B-AS1* were the dominant enhancer targets, with E1–E5, E8–E9, and E11–E12 exerting broad regulatory influence (**Figure S3G**). *MTAP* expression was modulated by E3–E9 and E11 in both cell types, suggesting partially conserved enhancer logic (**Figure S3F-3G**)

Independent RT-qPCR validation confirmed key E2G connections, including downregulation of *MTAP* and *CDKN2B* expression by enhancers E3, E5, and E9 in vascular fibroblast (**Figure S5A-S5B**), and regulation of *MTAP*, *CDKN2A*, *CDKN2B-AS1*, and *DMRTA1* by enhancers E2, E3, E5, and E9 in vascular smooth muscle cells (**Figure S5C-S5F**).

Together, these results demonstrate that the *9p21.3* risk haplotype contains multiple independently active enhancers with distinct cell-type–specific target repertoires, and nominate *MTAP* along with other previously predicted *9p21.3* cis-expressing genes as a consistently regulated effector gene across vascular fibroblast and smooth muscle cell contexts by multiple enhancers.

### High-Resolution Genetic and Engineered Fine-Mapping of *9p21.3* Variants Identifies MTAP as a CAD Effector Gene

Having established an atlas of enhancer-gene connectivities in vascular fibroblasts and vascular smooth muscle cells, we next performed systematic fine-mapping of the locus to identify which candidate genes represent the most likely effector targets of *9p21.3* genetic variation. Unlike other approaches that rely on predicted regulatory elements or sparse variant sampling, our dense 50-kb CRISPRi-MAC-Seq tiling design (**Figure 2C**) enabled unbiased high-resolution testing and dissection across the entire *9p21.3* locus (chr9:22081398-22125504) – spanning enhancers and *9p21.3* CAD variants.

To identify variant-flanking regions with the strongest gene-regulatory effects, we applied a sliding window approach that aggregates sgRNA transcriptional effects across the locus. Using 1,000-bp windows with 500-bp off-sets, we identified regulatory regions through permutation testing and integrated these with all 50 CAD haplotype variants (LD r^2^>0.8, 1000G) ^38^. This strategy confirmed and extended enhancer–gene connections identified earlier, revealing *MTAP*, *DMRTA1,* and *CDKN2B* as targets in vascular fibroblasts and *MTAP, CDKN2A*, *MIR31HG,* and *CDKN2B* in vascular smooth muscle cells (**Figure 2E-2F**; **Table 2**, **Figure S6A-6B**).

**Table 2.** Sliding window permutation analysis from the CRISPRi-MAC-Seq locus screen, showing regions with significant repression of one or more potential 9p21.3 target genes. The significance level used is a permutation z-score of = −1.96. Guides directly overlapping a ChromHMM enhancer annotation are indicated, as are those in an enhancer flanking region. SNPs that fall within a sliding window, and their status as constraint or credible set variants, are also indicated.

| celltype | coords | expression | n_guides | prm_mean | prm_sd | zscore | p | gene | enhancer | snp | cred set | constraint_overlap |
| --- | --- | --- | --- | --- | --- | --- | --- | --- | --- | --- | --- | --- |
| cfb | 22084000 | -0.601 | 8 | 0.011 | 0.287 | -2.133 | 0.016 | *DMRTA1* | E2 | rs1537370 |  |  |
| cfb | 22084500 | -0.683 | 5 | 0.003 | 0.333 | -2.06 | 0.02 | *DMRTA1* | E2 | rs1537370 |  |  |
| cfb | 22098500 | -0.63 | 8 | 0.014 | 0.306 | -2.108 | 0.017 | *DMRTA1* | E4 | rs4977574,rs2891168 |  |  |
| cfb | 22099500 | -1.016 | 3 | -0.005 | 0.475 | -2.129 | 0.017 | *MTAP* | E4 | rs1537371 | rs1537371 |  |
| cfb | 22105500 | -0.67 | 9 | -0.023 | 0.281 | -2.3 | 0.011 | *CDKN2B* | E5 flank | rs7859362 |  | rs7859362 |
| cfb | 22106000 | -0.569 | 13 | 0.024 | 0.221 | -2.68 | 0.004 | *DMRTA1* | E5 flank | rs7859362, rs10757275, rs6475609 |  | rs7859362 |
| cfb | 22110000 | -0.668 | 7 | -0.045 | 0.296 | -2.105 | 0.018 | *MTAP* | E7 | rs1412834 |  |  |
| cfb | 22116000 | -0.429 | 18 | -0.042 | 0.179 | -2.156 | 0.016 | *MTAP* | E9 | rs1004638,rs1537375,rs2383207,<br>rs1537374,rs1537376 |  | rs1537374, rs1537375,<br>rs1537376 |
| smc | 22086000 | -0.814 | 6 | -0.014 | 0.311 | -2.575 | 0.005 | *CDKN2A* | E3 | rs1970112 |  |  |
| smc | 22105500 | -0.629 | 9 | -0.013 | 0.275 | -2.243 | 0.012 | *MIR31HG* | E5 flank | rs7859362 |  | rs7859362 |
| smc | 22112000 | -0.397 | 23 | -0.044 | 0.168 | -2.104 | 0.018 | *MTAP* | E8 | rs7341791,rs7341786, |  |  |
| smc | 22112500 | -0.392 | 20 | 0.006 | 0.187 | -2.135 | 0.016 | *CDKN2B* | E8 | rs7341791,rs7341786,rs10511701 |  |  |
| smc | 22112500 | -0.452 | 20 | -0.048 | 0.177 | -2.285 | 0.011 | *MTAP* | E8 | rs7341791,rs7341786,rs10511701 |  |  |
| smc | 22116000 | -0.487 | 18 | -0.049 | 0.174 | -2.517 | 0.006 | *MTAP* | E9 | rs1004638,rs1537375,rs2383207,<br>rs1537374,rs1537376 |  | rs1537374, rs1537375,<br>rs1537376 |

In vascular fibroblasts, we identified significant regulatory effects of enhancers harboring CAD risk variants on *MTAP*, *DMRTA1*, and *CDKN2B* expression (**Figure 2E**; **Table 2**, **Figure S6A**). Specifically, enhancers E4, E7, and E9 significantly reduced *MTAP* transcript levels, while E2 (flanking rs2537370), E4 (rs4977574 and rs2891168), and E5 (containing rs7859362, rs10757275, and rs6475609) downregulated *DMRTA1*. Additionally, enhancer E5 significantly repressed *CDKN2B* expression. In vascular smooth muscle cells, we observed distinct but partially overlapping enhancer–gene relationships (**Figure 2F**; **Figure S6B**; **Table 2**). Enhancer E8 (containing rs7341791, rs7341786, and rs10511701) and enhancer E9 (harboring constraint variants rs1537374, rs1537375, and rs1537376) both significantly downregulated *MTAP* expression. Enhancer E8 also reduced *CDKN2B* levels. Furthermore, enhancer E5 (containing rs7859362) significantly repressed MIR31HG expression in smooth muscle cells, and modulated *CDKN2B* and *DMRTA1* in fibroblasts. We identified additional enhancer-to-gene (E2G) and SNP-to-gene (SNP2G) regulatory links within other enhancer regions of the *9p21.3* haplotype. Enhancers E3 (flanking rs1970122) significantly downregulated *CDKN2A* expression (**Figure S6A**; **Table 2**).

The resolution of GWAS associated genetic signals is inherently limited by the haplotype structure of the human genome, resulting in LD blocks of highly correlated variants. To nominate disease-driving variant(s) and effector gene(s) within the *9p21.3* locus, we integrated our sliding window CRISPRi tiling data with genetic constraint ^51^ and statistical fine-mapping metrics.

Our CRISPRi-MAC-seq tiling screen densely interrogated the 50-kb risk haplotype using 80 windows spanning the locus, 53 of which overlapped CAD-associated SNPs (median = 1, mean = 1.25, max = 8 guides per window) (**Table S8**). We first overlaid these data with a nucleotide level human constraint map derived from the Gnocchi model, generated from 76,156 whole-genome sequences from the Genome Aggregation Database (gnomAD)^51^. Gnocchi incorporates both local sequence context and genomic features to quantify the depletion of variation in tiled windows across the entire genome, indicating constraint regions undergoing natural selection. Although the *9p21.3* region lacks protein-coding genes, it exhibited a level of constraint (Z =1.10) comparable to coding sequences (Z=1.47), and markedly greater than the rest of the noncoding genome (Z=0.08; Wilcoxon P-value= 3.08×10^-12^) (**Figure 3A**).

**Figure 3.**
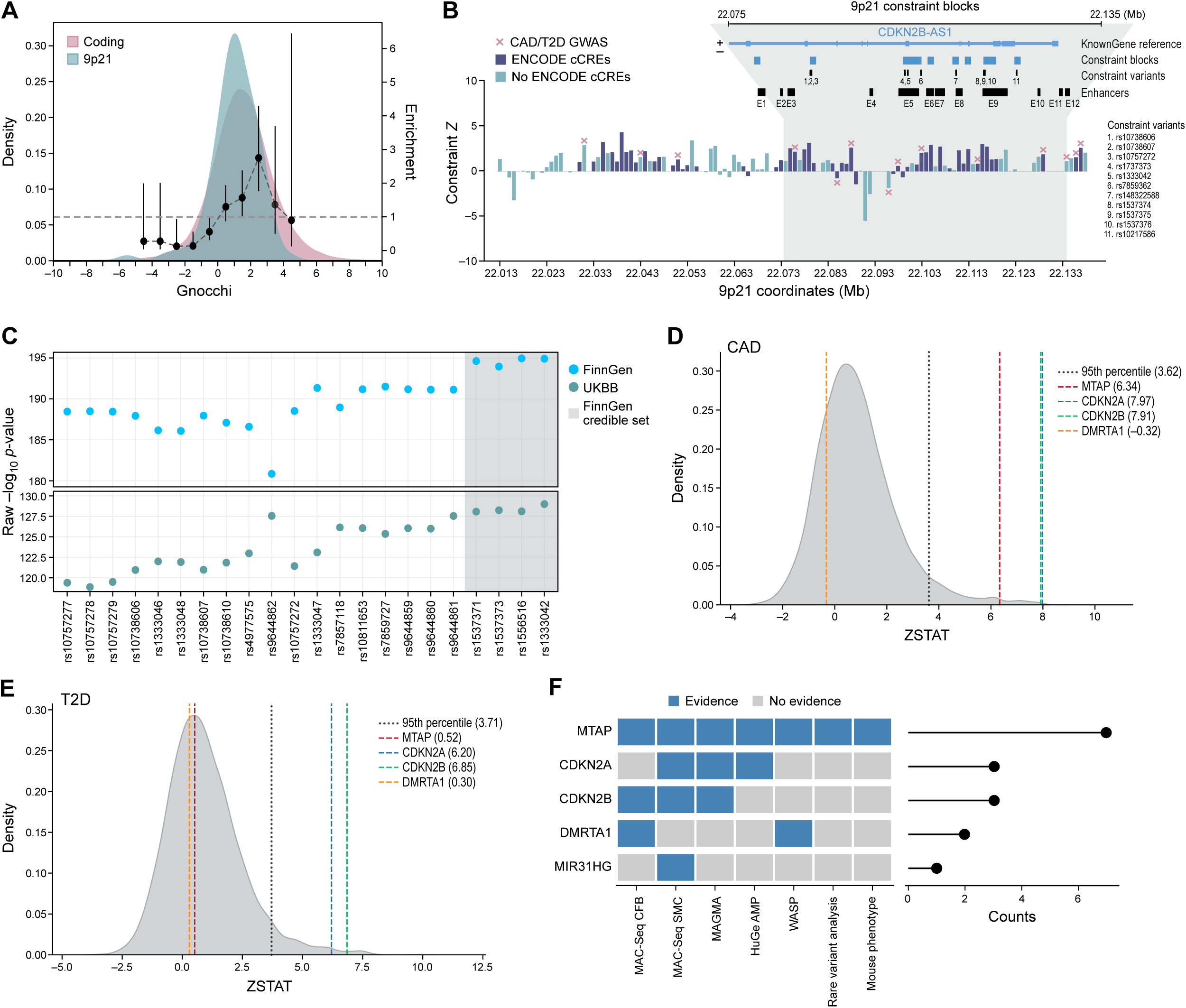
Convergent genomics evidence points to MTAP as the primary *9p21.3* CAD effector gene in vascular fibroblast and smooth muscle cells. (A) Density plot of Gnocchi scores across the genome. Windows overlapping coding regions (red color) compared to windows overlapping *9p21.3* non-coding regions (blue color). (B) Distribution of CAD/T2D variants in constraint regions spanning ∼120-kb region on the P-arm of chromosome 9. Overlaid on this plot is enhancer annotation for *9p21.3* locus along with constraint blocks and constraint variants flanking or overlapping enhancer region. (C) Dot plots showing fine-mapped variants associated with a combined trait for coronary revascularization, including coronary artery bypass grafting (CABG) and percutaneous coronary intervention (ANGIO) using data from large-scale population efforts from FinnGen and UKBB. Credible set variants are shaded grey and have the highest magnitude of association to CAD. (D-E) Using a MAGMA gene prioritization approach, we identified CAD-specific genes by calculating genome-wide ZSTAT scores for genes in comparison to a GWAS. Genes with ZSTAT scores exceeding the 0.95 percentile were considered significantly related to the GWAS trait. (D) First, we prioritized 9p21.3 genes against a CAD GWAS. The results indicated that MTAP, CDKN2A, and CDKN2B surpassed the significance cutoff. (E) To assess disease specificity, we then compared these results against a T2D GWAS. This analysis highlighted that MTAP’s association with CAD is disease-specific, as it did not meet the significance cutoff for T2D. (F) Converging evidence for CAD effector gene prioritization. The matrix denotes evidence to provide support (colored blue) for CAD driver genes across different modalities. Top scoring genes have the highest count across all evidence.

Notably, constraint signals were concentrated within E3-E4, E5, and E9 across both vascular fibroblasts and smooth muscle cells (**Figure 3B**, **Table S8**). The same region also harbored genetic markers identified by CAD GWAS ^51^. Regions containing CAD-associated variants showed significantly elevated constraint (median Z=1.57) relative to the non-associated background (median Z=0.08; Wilcoxon P-value=0.002), and were comparable to that observed across coding sequences (median Z=1.47; Wilcoxon P-value=0.95) (**Figure 3B**; **Table S8**).

We further derived credible set variants robustly associated with a combined trait for coronary revascularization, including coronary artery bypass grafting (CABG) and percutaneous coronary intervention (ANGIO), using data from large-scale biobanks including FinnGen and UKBB, resulting in the prioritization of four variants in the 95% credible set: rs1537371, rs1537373, rs1556516, and rs1333042 (**Figure 3C**; **Table 1**). Of these, only rs1537371 mapped to an enhancer (E4), exhibiting strong regulatory activity in fibroblast from our CRISPRi screen.

Integrating CRISPRi perturbation, constraint, and fine-mapping results, we found strong evidence that *MTAP* is the primary effector gene at the *9p21.3* locus. In vascular fibroblasts, three enhancers (E4, E7, E9) that harbor CAD-associated variants robustly downregulated *MTAP* expression (**Figure 2B**; **Table 2**). Notably, enhancer E9 contains three highly constrained variants (rs1537374, rs1537375, rs1537376), all of which significantly reduced *MTAP* expression when perturbed (Z-score=–2.16, P-value= 0.02; **Table 2**, **Table S8**). Furthermore, the enhancer window centered at chr9:22,099,500 (hg38), which harbors the fine-mapped variant rs1537371, elicited strong *MTAP* repression (Z-score=–2.13, P-value=0.02), implicating this SNP as a likely causal regulatory variant modulating *MTAP* expression in vascular fibroblasts. These three lines of evidence converging on the same target gene reveal MTAP as the primary target gene at the locus in vascular fibroblasts.

While the constraint signal for MTAP was strongest, we also identified constraint-based support for *DMRTA1* and *CDKN2B* in vascular fibroblasts. Specifically, the constrained variant rs7859362 flanking enhancer E5 was linked to reduced expression of both genes (**Table 2**, **Table S8**). In vascular smooth muscle cells, we found similar evidence implicating *MTAP*. The same set of constrained variants in enhancer E9 (rs1537374, rs1537375, rs1537376) also significantly downregulated *MTAP* in this cell type (Z-score=–2.52, P-value=0.006; **Figure 2D**; **Table 2**, **Table S8**), indicating *MTAP* as the most consistent and cell-type–spanning effector. Besides *MTAP*, the micro RNA *MIR31HG* was linked to the constrained variant rs7859362 flanking enhancer E5.

Altogether, these findings establish a convergent enhancer-variant-gene axis centered on *MTAP*, with E4 and E9 acting as regulatory hubs. Among all cis-genes evaluated, *MTAP*—located ∼269 kb from the tag SNP—is the only gene at the locus regulated by both fine-mapped and constrained variants in fibroblasts and smooth muscle cells. Although constrained variants support *DMRTA1*, *CDKN2B,* and *MIR31HG*, the integration of multiple orthogonal data sets identifies *MTAP* as the most consistently and robustly supported effector gene at the *9p21.3* CAD locus.

### *MTAP* Emerges as the Primary *9p21.3* CAD Effector Gene Through Convergent Genomic Evidence

Having identified *MTAP* as a genetically regulated target of multiple *9p21.3* enhancers and likely functional variants in vascular cells, we next sought further support for its candidacy as a CAD effector gene compared to other cis-expressing *9p21.3* genes through convergent genomic and population-scale evidence.

We first tested for allelic imbalance of *MTAP* expression at *9p21.3* in heterozygous donor-derived vascular cells using RNA-seq and array-based genotyping. A combined haplotype test (WASP analysis) ^52^ revealed significant allele-specific expression for *MTAP* (**Figure 3F**; **Table S9**), with lower expression from the CAD risk haplotype compared to the non-risk haplotype. Of all genes assessed at the locus, only *DMRTA1* additionally showed an allelic imbalance, also favoring lower expression from the risk haplotype. These data provide orthogonal validation that genetic variation at *9p21.3* impacts *MTAP* expression in primary vascular cells.

Given the significant allelic imbalance of *MTAP* expression observed in aggregated vascular cell datasets, we next assessed how genetic variation at the *9p21.3* haplotype affects cell type–specific chromatin accessibility and transcription at single-cell resolution. To this end, we analyzed single-cell Multiome data capturing both chromatin accessibility and gene expression profiles from human left anterior descending (LAD) coronary arteries of 23 patients (**Table S30**). After data integration and cell type annotation (see Methods), we identified 13 distinct vascular and stromal cell populations from a total of 107,825 single cells (**Figure S11A**). Using SCENT (single cell enhancer target gene mapping) ^53^, which quantifies associations between chromatin accessibility and gene expression at single-cell resolution. Focusing on the *9p21.3* region (chr9:22,075,000-22,135,000), we stratified cells by rs1537371 genotype—CC (homozygous non-risk), CA (heterozygous), and AA (homozygous risk)—within annotated (based on cell markers) vascular smooth muscle cells and fibroblasts (**Figure S11B**). This analysis identified a total 172 significant tile-to-gene pairs (P-value<0.05) across genotypes, with the number of regulatory connections varying by both genotype and cell type (**Figure S11C**; **Table S31**). In vascular fibroblasts, we observed a modest allele-dependent reduction of *MTAP* expression associated with rs1537371 genotype (CC vs CA; MAST P-value=0.009, FC=0.1050, **Figure S11D-S11E**, **Table S33**), aligning with reduced chromatin accessibility at the same locus. In smooth muscle cells, we did not observe any allele-dependent reduction in *MTAP* pseudobulk expression (**Figure S11F-S11G**; **Table S33**). Collectively, our analysis suggests variation at rs1537371 impacts *MTAP* levels in vascular fibroblast but not in smooth muscle cells, which is consistent with our CRISPRi-MAC-seq data (**Figure 2E**).

Next, we queried the publicly available Human Genetic Evidence Calculator (HuGE) ^54^, which integrates multiple human genetic evidence from rare and common genetic variation to quantify genetic support for involvement of genes linked to disease. *MTAP* demonstrated robust genetic support for CAD and related cardiovascular traits, with very strong associations for blood pressure (HuGE scores: SBP=83.89, DBP=56.10) and hypertension (HuGE=45.00) and significant associations with CAD (HuGE=20.00) and myocardial infarction (HuGE=20.00) (**Figure 3F**; **Figure S7A**; **Table S10**).

Furthermore, we assessed evidence from rare coding variation and CAD association for any of the candidate protein-coding genes in the *9p21.3* locus – *CDKN2A, CDKN2B, DMRTA1,* and *MTAP* using exome-sequencing data from four large cohorts namely the ExSeq study (N=57,178) and WGSeq study (N=6,809) from the Myocardial Infarction Genetics (40% European, 2% East Asian, 49% South Asian, and 7% African) and the UK Biobank 13K and 200K studies (European ancestry) ^55^. We previously reported the main finding associated with CAD ^56^, which incorporates loss-of-function prediction (LOFTEE) ^57^, splicing impact (SpliceAI) ^58^, ClinVar annotations ^59^, and very rare missense variants (allele frequency <0.01%) predicted to be damaging using our previously introduced method (pLoF+missense) ^56^. We observed the strongest nominal signal for MTAP (β= 0.6) among protein-coding genes, although not reaching statistical significance (P-value=0.09) (**Figure 3F**, **Table S11**). A negligible rare variant burden was detected for *CDKN2A*, *CDKN2B*, and *DMRTA1*.

Additionally, we explored evidence from *in vivo* genetic perturbation in mice. To assess phenotypic consequences of *MTAP* perturbation *in vivo*, we evaluated data from the International Mouse Phenotyping Consortium (IMPC) database ^60^, which catalogs the organ-system implication of all mammalian coding genes. Among all the *9p21.3* candidate target genes identified from our screen, *Mtap* haploinsufficiency was identified to be associated with abnormal vitellin vasculature morphology phenotype amongst other phenotypes – underscoring the role of *MTAP* in vascular disease development processes (**Figure 3F**, **Table S12**). We note that homozygous *Mtap* knockout was embryonic lethal with complete penetrance, displaying early developmental defects at E9.5. No data was reported for comparable CAD related phenotypes for other *9p21.3* associated genes, including *Cdkn2a*, *Cdkn2b*, or *Dmrta1* knockout mice (https://www.mousephenotype.org/).

To quantify the association of *9p21.3* cis-expressing genes to overall contribution to CAD trait, we performed a gene-level MAGMA ^61^ analysis using summary statistics from a recent GWAS meta-analysis (210,842 CAD cases among 1,378,170 participants) ^62^. Our data suggests that *MTAP* along with previously predicted *9p21.3* genes–*CDKN2A* and *CDKN2B–*ranked in the 99th percentile genome-wide (ZSTAT MTAP=6.34, CDKN2A=7.97, CDKN2B=7.91), exceeding the 95th percentile cut-off (**Figure 3D-3F**, **Table S13**). *DMRTA1* ranked below cut-off (DMRTA1=-0.32), and *MIR31HG* and *CDKN2B-AS1* were excluded in MAGMAs pipeline. Intriguingly, *MTAP*’s gene level contribution was disease-specific: among the genes exceeding the threshold for CAD, *MTAP* did not exceed the 95th percentile in a CAD-related control analysis on T2D GWAS summary statistics ^63^ (**Figure 3F**, **Table S32**). For schizophrenia ^64^ as a negative control, none of the genes exceeded the 95th percentile threshold (**Figure S8A**; **Table S14**).

To probe the translational relevance of *MTAP* in cardiovascular diseases, we investigated large-scale transcriptomic and proteomic datasets. Assessment of single-nucleus RNA-seq from human hearts^65^—including non-failing (NF), hypertrophic cardiomyopathy (HCM), and dilated cardiomyopathy (DCM) samples—suggests that compared to other *9p21.3* genes, *MTAP* showed the highest expression levels across all cardiac cell types (**Figure S7B**). Interestingly, compared to non-failing hearts, pseudobulk *MTAP* expression was significantly increased in HCM (P-value=0.027) and DCM (P-value=0.044) hearts (**Figure S7C**). At single cell level, we observed these significant increases in vascular cells isolated from HCM and DCM heart including in vascular smooth muscle cells, fibroblasts, among other cell types – suggesting dysregulation of MTAP expression in cardiovascular disease processes (**Figure S7D**). Additionally, analysis from the STARNET study profiling expressions in five CAD-relevant tissues ^66^, demonstrated consistent differential *MTAP* expression in CAD compared to the control cases (**Figure S7E**). Lastly, proteomic data from FinnGen ^30^ shows a significant increase in plasma MTAP levels in CAD patients compared to non-CAD patient controls (P-value=0.038; **Figure S7F**). Collectively, these findings implicate *MTAP* as a major effector gene at the *9p21.3* mediating CAD risk (**Figure 3F**).

### *MTAP* Regulates Transcriptional Programs That Orchestrate Vascular Development and Remodeling

Having identified *MTAP* as a genetically anchored CAD effector gene at the *9p21.3* locus, we next investigated how *MTAP* perturbation affects transcriptional networks involved in vascular cell function and disease susceptibility. *MTAP* encodes a metabolic enzyme that catalyzes the phosphorolysis of methylthioadenosine (MTA) to adenine and 5-methylthioribose-1-phosphate, playing essential roles in methionine salvage, polyamine metabolism, and cellular methylation capacity ^67^. The mechanistic role of *MTAP* in vascular pathophysiology remains largely unknown.

To dissect the downstream effects of *MTAP* loss, we employed an siRNA-mediated loss-of-function approach coupled with bulk RNA-seq, and profiled transcriptome-wide changes in primary human vascular fibroblasts and smooth muscle cells. We confirmed robust siRNA-mediated silencing of *MTAP* in both vascular fibroblasts and smooth muscle cells (KD efficiencies > 90%, **Figure S8B-C**), as confirmed by qPCR.

In vascular smooth muscle cells, *MTAP* knockdown elicited a subtle overall transcriptional response, with a total of 67 differentially expressed genes (DEGs; log₂FC>2; log₂FC<-2; P-adj<0.05), including 35 upregulated and 32 downregulated transcripts (**Figure 4A**; **Table S15**). Interestingly, *MTAP* knockdown significantly upregulated *CDKN2B* (log₂FC=4.9, P-adj=0.006), suggesting that *MTAP* may function as a negative regulator of cell cycle checkpoint pathways (**Table S15**, **Figure 4A**). Additionally, we identified several key genes that have been established to play key roles in vascular development and cardiovascular disease processes, functioning in extracellular matrix remodelling (Matrix Metalloproteinases - *MMP9*; log₂FC= −8.0; P-value= 5.82x 10^-05^) ^68^, elastic fiber assembly (Elastin microfibril interfacer 1 - *EMILIN1*; log₂FC= −6.6; P-value= 4.34 x 10^-06^) ^69^, arterial specification (Neurogenic locus notch homolog protein 3 - *NOTCH3*; log₂FC= 6.7; P-value=4.38x 10^-05^) ^70^, smooth muscle cell proliferation (Platelet derived growth factor receptor - *PDGFRA*; log₂FC= 5.1; P-value= 6.88x 10^-07^) ^71,72^, and development and progression of atherosclerosis (Cathepsin S - *CTSS*; log₂FC= −26.1; P-value= 2.36x 10^-17^) ^71^ – suggesting that *MTAP* plays a key role in regulating processes crucial to vascular diseases.

**Figure 4.**
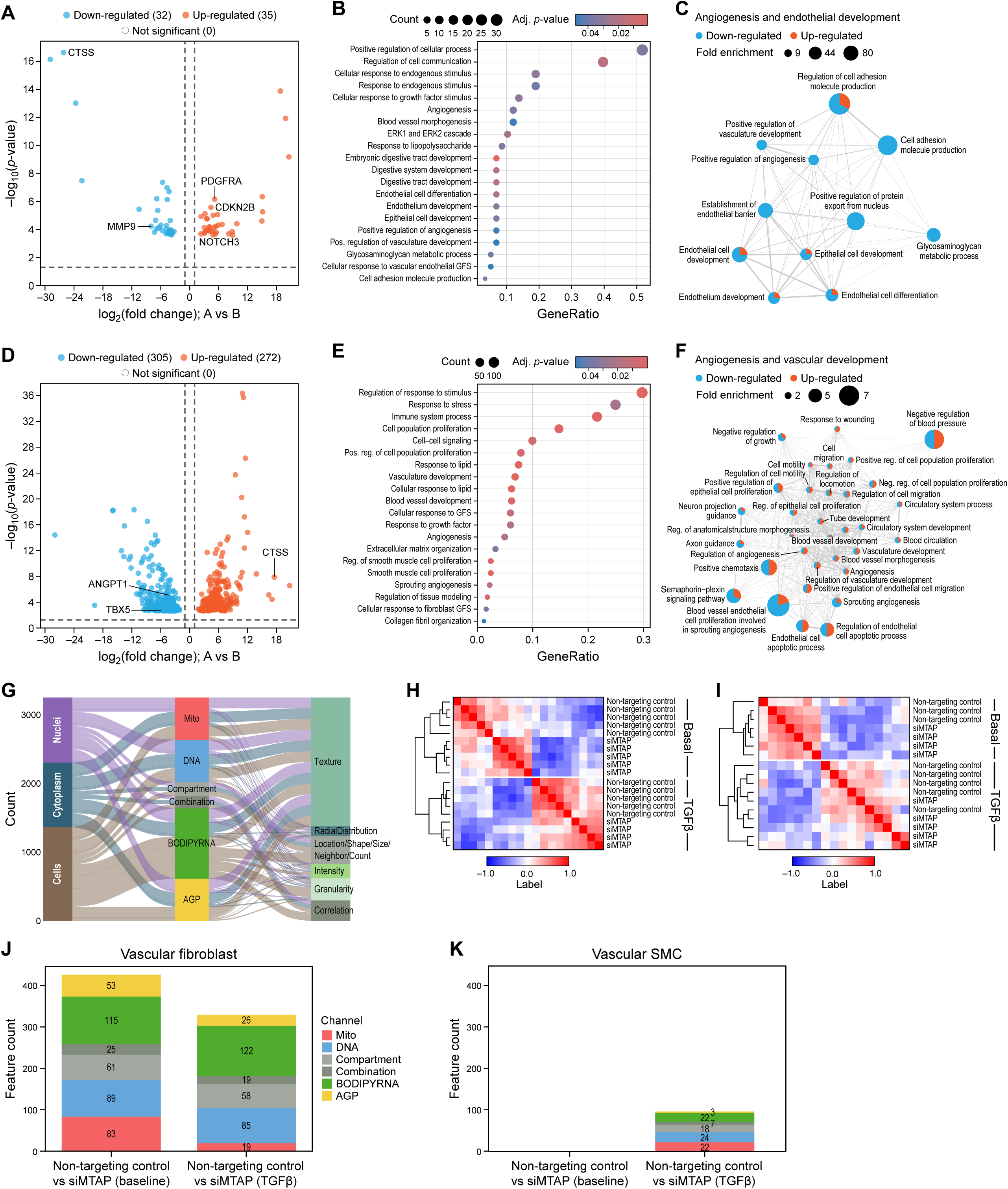
MTAP regulates transcriptional programs that orchestrate vascular development and remodeling (A) Volcano plots showing differentially expressed genes in vascular smooth muscle cells as a result of MTAP knockdown via siRNA. Genes with a P< 0.05 are considered significant. (B) Gene ontology (GO) analysis showing biological processes associated with DEGs (p-adjusted<0.05) as a result of MTAP suppression in vascular smooth muscle cells. (C) Network analysis of gene enrichment results to find relevant functional modules. Resulting clusters represent cellular processes overrepresented in statistically significant differentially expressed genes as a result of MTAP knockdown in vascular smooth muscle cells. Within each pathway in a cluster, we can visualize genes as upregulated (red) or downregulated (blue) based on their annotation to that pathway. The size of each pathway denotes its fold enrichment, indicating how well that pathway is enriched in our differentially expressed genes compared to the background. (D) Volcano plots showing differentially expressed genes in vascular fibroblasts as a result of MTAP knockdown via siRNA. Genes with a P< 0.05 are considered significant. (E) Gene ontology (GO) analysis showing biological processes associated with DEGs (p-adjusted<0.05) as a result of MTAP suppression in vascular fibroblasts. (F) Network analysis of gene enrichment results to find relevant functional modules. Resulting clusters represent cellular processes overrepresented in statistically significant differentially expressed genes as a result of MTAP knockdown in vascular fibroblasts. Within each pathway in a cluster, we can visualize genes as upregulated (red) or downregulated (blue) based on their annotation to that pathway. The size of each pathway denotes its fold enrichment, indicating how well that pathway is enriched in our differentially expressed genes compared to the background. (G) Alluvial plot showing total number of extracted LipocyteProfiler features from vascular fibroblast and smooth muscle cells by cellular compartments, stained organelle/structure, and feature attribute (object size, feature intersection, texture, and others). (H) Pearson correlation matrix of full extracted feature profiles from basal groups (siNTCtr and siMTAP) as well as in TGF-β treated groups (siNTCtr and siMTAP) in vascular fibroblast. (I) Pearson correlation matrix of full extracted feature profiles from basal groups (siNTCtr and siMTAP) as well as in TGF-β treated groups (siNTCtr and siMTAP) in vascular smooth muscle cells (J) Stacked bar graph showing counts of differentially changed LipocyteProfiler features at baseline (siNTCtr vs siMTAP) and in response to TGF-β (siNTCtr vs siMTAP) in vascular fibroblasts. All features with q-values ≤ 0.2 were considered significant. (K) Stacked bar graph showing counts of differentially changed LipocyteProfiler features at baseline (siNTCtr vs siMTAP) and in response to (TGF-β siNTCtr vs siMTAP) in vascular smooth muscle cells. No differentially changed features were observed at baseline in vascular smooth muscle cells. All features with q-values ≤ 0.2 were considered significant.

Gene Ontology (GO) enrichment analysis of DEGs revealed that *MTAP* loss in smooth muscle cells affects transcriptional programs related to angiogenesis (GO:0045766; P-value=0.046; fold enrichment=8.8), vasculature development (GO:1904018; P-value=0.048, fold enrichment = 8.6), and response to growth factor (GO:0070848; P-value=0.04, fold enrichment=4.3), among others (**Figure 4B**; **Table S16**). To identify higher-order functional relationships among enriched biological processes and identify coordinated transcriptional modules, we constructed a GO network based on shared gene content (see Methods). Four major clusters emerged, including ‘Signal transduction in development’, ‘Vascular and digestive system development’, ‘TLR-mediated immune response’, and ‘Endothelial development and angiogenesis’ (**Figure 4C**; **Table S17**; **Figure S9A**). The module ‘Endothelial development and angiogenesis’ demonstrated the highest average fold enrichment (mean=26.52, range=80.14 to 8.56) and contained core angiogenic processes which are coordinatively down-regulated, including “positive regulation of angiogenesis” (GO:0045766; P=0.046, fold enrichment = 8.83), “endothelial cell development” (GO:0001885; P-value=0.016, fold enrichment=23.87), and “vasculature development” (GO:1904018; P-value=0.048, fold enrichment=8.56) (**Table S16**). Angiogenesis-related processes showed the strongest enrichment (highest fold change), indicating an induction of pro-angiogenic signaling (**Figure 4C**; **Table S17**). These results indicate a role for *MTAP* in activation of a comprehensive pro-angiogenic transcriptional program in vascular smooth muscle cells.

In contrast to vascular smooth muscle cells, primary human vascular fibroblasts exhibited a substantially broader transcriptional response to *MTAP* perturbation, with 577 genes reaching statistical significance (DEGs; log₂FC>1; log₂FC<-2; padj<0.05). This extensive perturbation response comprised 272 upregulated and 305 downregulated transcripts (**Figure 4D**; **Table S18**). Pathway analysis revealed enrichment of a broader spectrum of critical vascular-relevant biological processes following *MTAP* knockdown, including cellular response to fibroblast growth factor stimulus (GO:0044344; P-value=0.037, fold enrichment=3.4), extracellular matrix organization (GO:0030198; P-value=0.034, fold enrichment=2.3), response to lipid (GO:0071396, P-value= 0.0005, fold enrichment= 2.5), regulation of cell proliferation (GO:0002043, P-value=0.013, fold enrichment=7.4), blood vessel development (GO:0001568; P-value=0.005, fold enrichment = 2.1), and vasculature development (GO:0001944; P-value=0.002, fold enrichment=2.2), among many others (**Figure 4E**; **Table S19**). Network clustering of DEGs in primary fibroblasts revealed substantially greater complexity and higher degrees of interconnectivity compared to vascular smooth muscle cells, with seven major functional modules (modularity=0.15) reflecting the broader transcriptional response observed in fibroblasts (**Figure 4F**, **Figure S9B**, **Table S19**). One of the modules with a high degree of node connectivity (n=41 pathways, average connectivity degree = 308.36, **Table S20**) and high intracluster betweenness indicating robust pathway functionality within the module, “Angiogenesis and vascular development”, emerged as a significantly enriched cluster, containing 146 genes with an average fold enrichment of 2.75 (range = 1.66 to 7.35, P-adj < 0.05). This module encompassed core vascular processes including “blood vessel development” (GO:0001568; P-value=0.005, fold enrichment = 2.1), “angiogenesis” (GO:0001525; P-value=0.02, fold enrichment = 2.009), “regulation of vasculature development” (GO:1901342; P-value=0.02, fold enrichment = 2.32), and “regulation of cell population proliferation” (GO:008284; P-value=0.005, fold enrichment = 1.89) (**Figure 4F**; **Table S19**). Within the “Angiogenesis and vascular development” module, hub pathways with highest connectivities included Angiopoeitin 1 (*ANGPT1)* (log₂FC=-8.34, P-value=6.2 x 10^-06^), T-box 5 (*TBX5)* (log₂FC=-8.61, P-value=0.001) – both of which are crucial in vascular development and cardiac morphogenesis ^73,74^ (**Figure 4D-F**; **Table S20**). Additional modules in vascular fibroblasts included: “inflammatory and stress response” (n = 55 pathways, average fold enrichment=3.3), containing cytokine signaling and stress response pathways; “MAPK and phosphorylation signaling” (n=15 pathways, average fold enrichment=1.74), encompassing kinase cascades and post-translational regulation; “immune cell activation and migration” (n=38 pathways, average fold enrichment=3.97), representing chemotaxis and immune cell recruitment processes; “cell differentiation and state transition” (n=68 pathways, average fold enrichment=6.94), containing broader developmental programs; and “lipid metabolism and blood pressure regulation” (n=95 pathways, average fold enrichment=7.56) (**Figure S9B**; **Table S20**). These findings suggest that *MTAP* regulates multiple vascular disease-relevant pathways across diverse biological axes, including inflammation, angiogenesis, ECM remodeling, and metabolic homeostasis.

Together, the transcriptional programs activated by *MTAP* perturbation in primary vascular fibroblasts closely mirror those observed in pathological vascular remodeling, including atherosclerosis, aneurism, and peripheral artery Disease (PAD). The coordinated regulation of angiogenesis-related processes, vascular development, inflammation and metabolic homeostasis pathways suggest that reduced *MTAP* expression may promote a pro-atherogenic cellular phenotype.

### *MTAP* Functions as a Critical Regulator of Pathological Vascular Cell-State Transitions

Given the transcriptional programs modulated by *MTAP* suppression—particularly those linked to angiogenesis, ECM remodeling, and vascular development—we next investigated whether *MTAP* directly modulates disease-relevant cellular phenotypes. During atherosclerosis development, resident vascular wall cells undergo pathological state transitions characterized by altered contractile properties, enhanced proliferative capacity, and increased extracellular matrix production ^75^. A hallmark of vascular pathology is therefore the transition of quiescent vascular fibroblasts and smooth muscle cells into activated, disease-like phenotypic states. This phenotypic switch is driven by pro-fibrotic cues, most notably transforming growth factor β (TGF-β), and is characterized by morphological changes, upregulation of α-smooth muscle actin (αSMA), and enhanced matrix production, including collagen I ^76,77^.

To assess whether *MTAP* functions as a regulator of these transitions, we employed cardiometabolic-disease oriented high-content imaging in primary human vascular fibroblasts and smooth muscle cells following *MTAP* knockdown, under both basal and TGF-β-stimulated conditions.

We first applied LipocyteProfiler ^78^, a high-dimensional image-based profiling platform that quantifies 3,252 morphological and cellular features across subcellular compartments—including nucleus (Hoechst), mitochondria (MitoTracker Red), actin cytoskeleton and plasma membrane (phalloidin, wheat germ agglutinin), lipid droplets, nucleoli, and cytoplasmic RNA (BODIPY and SYTO14)—using multiplexed fluorescent staining (**Figure 4G**; **Table S21**). These features quantify morphological changes based on intensity, texture, granularity, and spatial organization across each subcellular compartment, providing comprehensive phenotypic characterization following *MTAP* depletion (**Figure 4G**) ^78^.

Hierarchical clustering of LipocyteProfiler features revealed distinct grouping patterns across experimental conditions (control, siMTAP, TGF-β, and TGF-β+siMTAP) in fibroblasts (**Figure 4H**, **Table S22**) and smooth muscle cells (**Figure 4I**, **Table S23**), demonstrating that the assay robustly captures both basal and stimulus-responsive morphological states. Separation between siMTAP and non-targeting control was more pronounced in vascular fibroblasts than smooth muscle cells (**Figure 4H-I**).

In primary vascular fibroblasts, *MTAP* knockdown resulted in substantial morphological remodeling under both basal conditions (n=426; q-value≤0.2) and in response to TGF-β exposure (n=329 features; q-value ≤ 0.2) (**Figure 4J**; **Table S24**). These changes spanned multiple cellular compartments, with strong effects observed in AreaShape, Texture, and Radial Distribution following *MTAP* depletion—indicative of shift in cytoskeletal dynamics, subcellular organization, and cell shape (**Figure 4J**; **Table S24**). Notably, features derived from the AGP (actin-Golgi-plasma membrane) channel showed consistent and coordinated alterations across both basal and stimulated conditions upon *MTAP* knockdown, suggesting that *MTAP* impacts structural reprogramming across the cytoskeleton and organelles (**Figure 4J**; **Table S24**). These findings suggest that *MTAP* perturbation results in profound morphological changes in both basal and TGF-β conditions, with importance for mitochondrial dynamics and cytoskeletal organization during cellular activation.

Vascular smooth muscle cells showed minimal response to *MTAP* perturbation under basal conditions and a modest morphological response under TGF-β stimulation (n = 96 features; q-value ≤ 0.2) (**Figure 4K**, **Table S25**). The differential magnitude of morphological responses between cell types (vascular fibroblasts showing 5.8-fold more affected features than smooth muscle cells following *MTAP* knockdown)–along with more pronounced transcriptome-wide changes following *MTAP* knockdown in vascular fibroblasts compared to smooth muscle cells–suggests cell type-specific roles for *MTAP* in regulating cellular responses to pro-fibrotic stimuli and supports broader effects on fibroblast cellular programs.

To specifically assess *MTAP’*s role in pathological cell state transitions relevant to vascular disease, we quantified alpha smooth muscle actin (αSMA) fiber formation—a hallmark of the transition from quiescent fibroblasts and contractile smooth muscle cells to activated, contractile disease-relevant phenotypes which are increasingly recognized as key contributors to CAD. This cellular reprogramming in response to TGF-β and other growth factors and inflammatory factors is characterized by acquisition of contractile stress fiber networks and enhanced matrix-producing capacity, and represents a convergent pathological endpoint across multiple vascular cell types during atherosclerosis and arterial remodeling ^76,77^. αSMA fiber area was quantified using automated high-content imaging analysis ^65^ with immunofluorescent αSMA staining (intensity threshold > 1500 arbitrary units), with measurements normalized to total cell number per field to control for proliferation effects.

In primary vascular fibroblasts, TGF-β stimulation of control cells (scrambled siRNA) significantly increased αSMA fiber area compared to basal conditions (12.75-fold increase, P-value< 0.0001), confirming robust disease-relevant fibroblast activation (**Figure 5A**). Strikingly, *MTAP* knockdown significantly increased this TGF-β-induced vascular state transitioning, with TGF-β-stimulated siMTAP treated cells showing 3.84-fold greater αSMA fiber area compared to TGF-β-stimulated controls (P-value=0.003), and a 30.5-fold greater αSMA fiber area compared to siMTAP at basal conditions (**Figure 5A**). Even in the absence of TGF-β, *MTAP*-depleted fibroblasts showed modestly elevated αSMA fiber area compared to controls (1.6-fold, P-value=0.025), suggesting that *MTAP* restrains spontaneous and induced contractile reprogramming.

**Figure 5.**
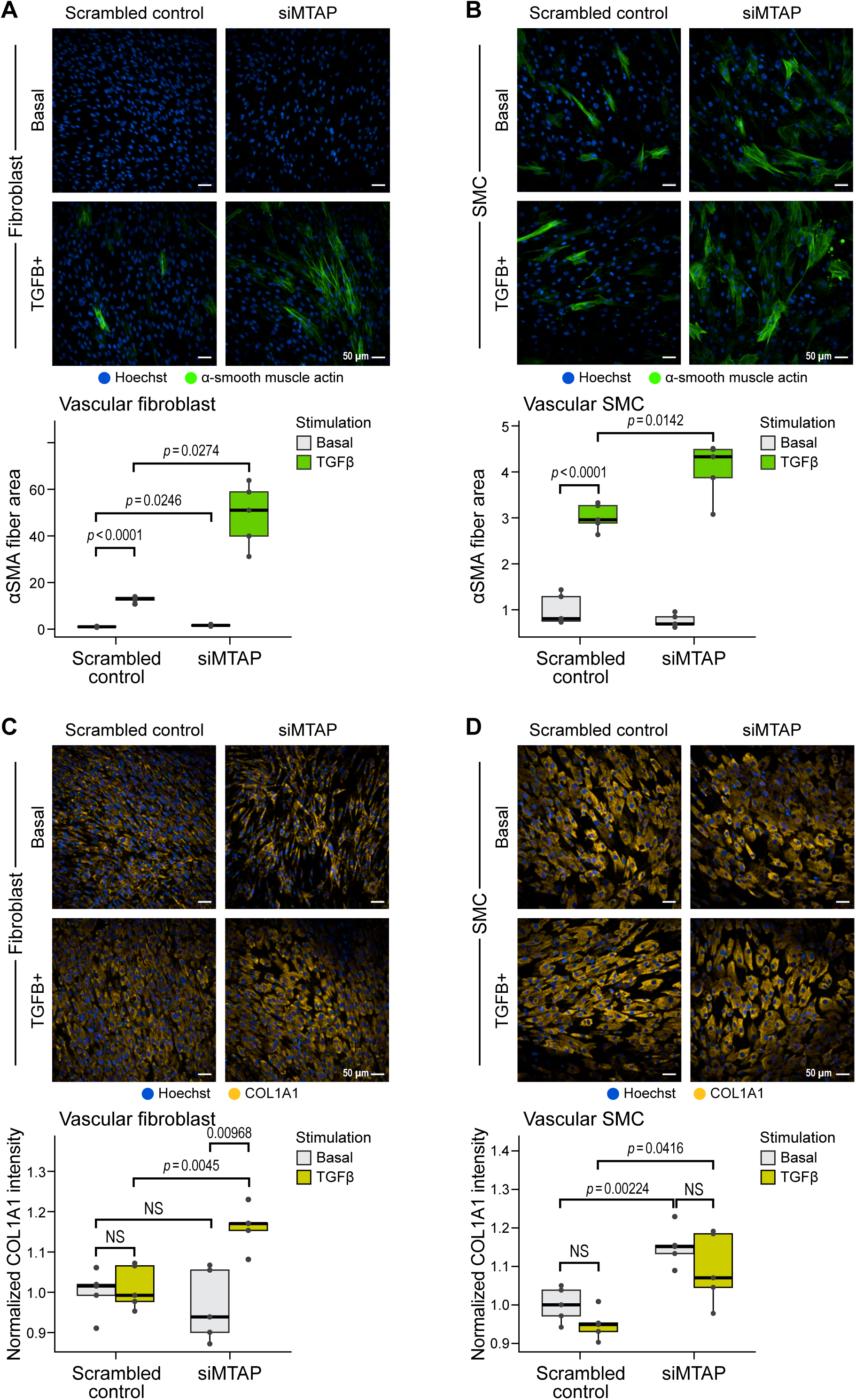
*MTAP* functions as a critical regulator of pathological vascular cell transitions (A) Representative images and box plots showing impact of siRNA mediated MTAP knockdown on alpha smooth muscle actin fiber areas (αSMA) at baseline and in response to TGF-β treatment in vascular fibroblasts. One-way ANOVA was to assess statistical significance followed by pairwise t-tests with Bonferroni correction. Scale bar (bottom right) = 50 µM. (B) Representative images and box plots showing impact of siRNA mediated MTAP knockdown on alpha smooth muscle actin fiber areas (αSMA) at baseline and in response to TGF-β treatment in vascular smooth muscle cells. One-way ANOVA was to assess statistical significance followed by pairwise t-tests with Bonferroni correction. Scale bar (bottom right) = 50 µM. (C) Representative images and box plots showing impact of siRNA mediated MTAP knockdown on Col1a1 levels (Col1a1) at baseline and in response to TGF-β treatment in vascular fibroblasts. One-way ANOVA was to assess statistical significance followed by pairwise t-tests with Bonferroni correction. Scale bar (bottom right) = 50 µM. (D) Representative images and box plots showing impact of siRNA mediated MTAP knockdown on Col1a1 levels (Col1a1) at baseline and in response to TGF-β treatment in vascular smooth muscle cells. One-way ANOVA was to assess statistical significance followed by pairwise t-tests with Bonferroni correction. Scale bar (bottom right) = 50 µM.

These findings establish *MTAP* as a critical negative regulator of the fibroblast-to-myoblast phenotypic switch that partially drives vascular fibrosis and arterial remodeling, consistent with our LipocyteProfiler high-dimensional imaging data demonstrating extensive changes in actin-Golgi-plasma membrane (AGP) channel features upon *MTAP* deletion and TGF-β stimulation—morphological signatures characteristic of pathological vascular cell state transitions.

In primary vascular smooth muscle cells, which constitutively express αSMA, TGF-β-stimulation of control cells resulted in enhanced αSMA fiber organization and intensity (3.0-fold increase in fiber area, P-value<0.0001) (**Figure 5B**). *MTAP* knockdown further augmented this effect (1.35-fold increase, P-value=0.014). Under basal conditions, no significant differences were observed between control and *MTAP*-depleted smooth muscle cells, indicating that *MTAP* modulates inducible rather than constitutive αSMA architecture in this cell type.

Given our transcriptomic evidence for *MTAP* regulation of extracellular matrix organization pathways (**Figure 4E**, **Table S19**), we next assessed whether *MTAP* perturbation affects collagen I production—a key effector mechanism in vascular fibrosis and atherosclerotic plaque formation and a defining functional hallmark of the sustained, pathological matrix-producing phenotype that drives vascular fibrosis. While αSMA expression represents the early contractile reprogramming of the activated cell state switch, enhanced collagen I synthesis reflect the long-term commitment to the pathological phenotype responsible for arterial wall thickening, perivascular fibrosis, and atherosclerotic plaque remodeling ^79,80^. We quantified COL1A1 protein levels using automated immunofluorescence imaging with intensity measurements normalized to cell number.

In vascular fibroblasts under basal conditions, COL1A1 levels were comparable between control and *MTAP*-depleted cells, indicating minimal effects on constitutive collagen expression (**Figure 5C**). TGF-β-stimulation of control cells also did not lead to an increase in COL1A1 levels. However, *MTAP* knockdown largely sensitized cells to TGF-β stimulation as reflected in substantial upregulation of COL1A1 in TGF-β-treated siMTAP cells compared to both their basal state (1.2-fold increase, P=0.0097) and TGF-β-stimulated controls (1.15-fold increase, P-value=0.0045). This enhancement of TGF-β responsiveness demonstrates that *MTAP* restrains pro-fibrotic signaling in vascular fibroblasts, and functions as a critical molecular brake preventing the transition to the sustained matrix-producing activated fibroblast phenotype that characterizes pathological vascular remodeling.

In vascular smooth muscle cells, a distinct pattern emerged where *MTAP* depletion resulted in constitutively elevated COL1A1 levels under both basal (1.8-fold increase vs. control, P-value=0.002) and TGF-β-stimulated conditions (1.5-fold increase vs. TGF-β control, P-value=0.042) (**Figure 5C**). This constitutive elevation suggests that *MTAP* loss promotes a partial phenotypic shift in smooth muscle cells, characterized by enhanced baseline matrix synthesis capacity—a pathological feature associated with the synthetic smooth muscle cell phenotype that contributes to atherosclerotic plaque development and arterial stiffening. Unlike fibroblasts, smooth muscle cells did not show additional TGF-β-induced increases in COL1A1 following *MTAP* knockdown, suggesting that *MTAP* regulates the constitutive matrix-producing capacity rather than specifically modulating stimulus-responsive activation and matrix production in this cell type.

The convergent findings from high-dimensional morphological profiling and targeted functional assays reveal that *MTAP* functions as a multifaceted regulator of pathological vascular cell state transitions. The preferential effects of *MTAP* knockdown on mitochondrial morphology and cytoskeletal organization (detected by LipocyteProfiler) align with the enhanced contractile protein expression (αSMA) and extracellular matrix production (COL1A1) observed through targeted imaging, suggesting coordinated regulation of the cellular machinery required for pathological vascular cell activation.

Mechanistically, our data establishes a functional link between reduced *MTAP* expression and pro-atherogenic cellular remodeling. *MTAP* knockdown in primary vascular fibroblasts and smooth muscle cells induces transcriptional programs associated with TGF-β signaling, ECM remodeling, and and pathological activation of matrix-producing vascular cells, as well as increased αSMA fiber formation and collagen I production. These effects suggest that loss of *MTAP* function promotes pathological cell state transitions that underlie vascular fibrosis and disease progression.

## Discussion

The chromosome *9p21.3* locus is the most significant genetic risk locus for CAD. In many populations, the risk haplotype occurs in approximately 50% of individuals, and its additive effect on disease susceptibility accounts for an estimated 10-15% of all CAD cases in the United States, making it the single largest genomic contributor to national healthcare due to CAD-related morbidity and mortality ^21,81^. In a recent large-scale analysis from FinnGen, the minor allele of a sentinel SNP at this locus (rs7859727, in the CAD risk haplotype, **Table 1**) was shown to contribute 447 disability-adjusted life years (DALYs) per 100,000 individuals per year—again, the highest DALY burden of any variant genome-wide ^82^. Despite this outsize public health impact, the effector gene(s), relevant cell types, and variant–mechanism relationships at *9p21.3* have remained incompletely defined. Here, we construct the most comprehensive functional perturbation-phenotype map of the *9p21.3* locus in human vascular cells to date, integrating epigenomic and transcriptional profiling, dense CRISPR interference tiling, constraint and fine-mapping analyses, high-content phenotyping, and disease-relevant cell state transitions in primary human vascular fibroblasts and smooth muscle cells. Through this systematic dissection, we identify *MTAP* as a genetically-anchored effector gene at *9p21.3* and demonstrate that its perturbation promotes pathological cell-state transition relevant to CAD initiation and progression.

We first establish that the *9p21.3* risk haplotype forms a modular enhancer cluster, comprising 12 independent enhancers dynamically activated by inflammatory and metabolic stimuli. We found that this dense enhancer cluster does not represent a super-enhancer, but rather forms a modular enhancer hub in vascular wall cells capable of integrating genetic signals through vascular cell-specific regulatory programs. This finding is contrary to previous studies that suggest enhancer clusters at *9p21.3* locus contain super-enhancers in cancer cells ^83^, suggesting a disease-specific and cell-type specific enhancer behavior as it relates to *9p21.3* locus that have previously not been identified. Heritability partitioning analysis implicates stimulated fibroblasts and smooth muscle cells as important mediators of CAD genetic risk. This context-specific regulatory activity provides a mechanistic basis for prior epidemiological studies which suggest interactions between the *9p21.3* locus and aging ^16^, diet ^25^, hypercholesterolemia ^26^, and hyperglycemia ^44^ and type 2 diabetes ^27^ — while others suggest its effects are largely independent of traditional CAD risk pathways ^28^.

Our tiling CRISPRi-MAC-seq screen identified enhancer-gene (E2G) as well as variant-enhancer-gene (SNP2G) regulatory relationships in vascular fibroblasts and smooth muscle cells. These data revealed that the *9p21.3* haplotype contains multiple independently active enhancers with cell-type–specific target genes, regulating known genes (e.g., *CDKN2A*, *CDKN2B*) and newly prioritized candidates such as *MTAP*, *MIR31HG* and *DMRTA1*. Integration with fine-mapped variants (i.e., rs1537371) and constrained sequence elements (e.g., rs1537376, rs7859362) revealed a convergent regulatory axis centered on *MTAP*. Notably, rs1537371, a fine-mapped CAD risk variant window located in enhancer E4, significantly down-regulated MTAP in fibroblasts when perturbed. While this variant was previously linked to p14ARF and p16INK4a (*CDKN2A*) expression ^84^ and endothelial cell senescence via SABT2 and CUX1, our CRISPRi-MAC-seq and single cell Multiome analysis of vascular cells analysis highlight a pleiotropic regulatory role for rs1537371, acting through distinct target genes depending on cell type—*CDKN2A* in endothelial cells and *MTAP* in fibroblasts—underscoring the importance of cell-type–resolved functional mapping for understanding complex trait loci.

While several other *9p21.3* genes—*CDKN2A/B*, *DMRTA1*, *MIR31HG*—were affected by enhancer perturbation in a context-dependent manner, none showed the same integration of cross-cell-type regulation, fine-mapping support, and enhancer constraint as *MTAP*. These findings suggest that *MTAP* is the primary effector gene at the locus, with other genes possibly contributing contextually or secondarily.

*MTAP* encodes methylthioadenosine phosphorylase, a metabolic enzyme catalyzing a key step in methionine salvage and polyamine metabolism, with downstream effects on cellular methylation potential ^67^. MTAP loss is well studied in cancer ^85^, but its role in vascular pathophysiology has been largely unexplored. In mice carrying the human APO*E3 Leiden transgene, Mtap haploinsufficiency leads to a significantly increased aortic sinus lesion size following an atherogenic diet ^86^, suggesting a role in atherogenesis. However, the cellular mechanisms remained undefined. Here, we use a combination of siRNA mediated knockdown in primary human vascular cells with bulk RNA-seq, and high-dimensional phenotyping as well as targeted CAD relevant image-based assays to define MTAP’s role in vascular cell biology. MTAP depletion in fibroblasts and smooth muscle cells activates TGF-β-responsive transcriptional programs that mirror those observed in vascular remodeling and disease—including angiogenesis, ECM organization, and inflammation. These signatures overlap with those observed in atherosclerosis, aortic aneurysm, and peripheral artery disease, indicating that reduced *MTAP* expression may promote a pro-atherogenic cellular phenotype. In smooth muscle cells, MTAP depletion also enhances Toll-like receptor (TLR) signaling ^87^, a key driver of vascular inflammation and CAD progression. This raises the possibility that MTAP modulates immune-metabolic crosstalk in vascular cells, restraining TLR2 and TLR4 mediated pro-inflammatory signaling.

Notably, our study demonstrates that MTAP serves as a repressor of pathological cell state transitions in both vascular fibroblasts and smooth muscle cells. In fibroblasts, we find that MTAP knockdown induces increased sensitivity to TGF-β stimulation, lowering the threshold for activation of fibrotic transitions in the presence of pro-atherogenic stimuli. The involvement of MTAP in pathophysiological cell state transitions are further supported by high-dimensional image-based phenotyping using LipocyteProfiler, which shows broad structural remodeling across >400 features, most notably coordinated cytoskeletal and mitochondrial remodeling. Importantly, these unbiased phenotypes were validated by targeted disease-relevant immunofluorescence imaging, which revealed that MTAP knockdown increases αSMA fiber formation and collagen I production—hallmarks of pathological, activated cell states. Interestingly, recently, Salido et al. (2025) showed that enhancer perturbation in IPSC derived vascular smooth muscle cells activated an osteochondrogenic program.

The identification of MTAP as a negative regulator of pathological vascular cell state transitions also suggests potential therapeutic strategies. Small molecules that enhance MTAP activity or downstream methionine salvage pathway flux might antagonize pro-fibrotic signaling and reduce vascular disease progression, particularly in individuals carrying *9p21.3* risk variants. Importantly, the therapeutic relevance of MTAP may extend more broadly, offering benefit even in individuals without non-coding *9p21.3* risk alleles.

In summary, our functional characterization demonstrates that MTAP serves as a critical molecular brake on pathological vascular cell state transitions, with its perturbation promoting the activated, pro-fibrotic cellular phenotypes characteristic of atherosclerotic vascular diseases.

## Methods

### Data and Code Availability

Code used in all analyses will be made available and public on GitHub at https://github.com/MelinaClaussnitzerLab/V2F-9p21.3-CAD and https://github.com/Antidington/LAD_phase1_9p21. All data supporting findings from this study are included in the main figures of the manuscript as well as in the supplemental figures.

### Cell Lines and Media Condition

All primary cells used for epigenetic profiling of vascular cells were commercially sourced. We utilized primary human cells, including atrial fibroblast (Lonza, CC-2903), ventricular fibroblast (Lonza, CC-2904), coronary artery endothelial cells (Lonza, CC-2585), pulmonary artery smooth muscle cells (Lonza, CC-2581), coronary artery smooth muscle cells (Lonza, CC-2583), and adipose derived microvascular pericytes (Angioproteomie, cAP-0043).

For siRNA treatment and high-content imaging experiments, primary human vascular fibroblasts were purchased commercially as well. We utilized human vascular fibroblasts (Cell Systems, ACBRI 5118) and primary human coronary artery smooth muscle cells (ATCC; PCS-100-021). Herein, we describe these cells as vascular fibroblasts and vascular smooth muscle cells when reporting findings in the main text.

Cells used for immortalized vascular cell line generation were commercially purchased. We utilized immortalized human vascular fibroblast (Applied Biological Materials Inc. T0446) and human coronary artery smooth muscle cells (Applied Biological Materials Inc., T0557).

### Cell Maintainance

All fibroblast cell lines were cultured in Lonza FGM-3 (CC-4526) and all smooth muscle cell lines were cultured in Lonza SmGM-2 (CC-3182) unless otherwise noted. Endothelial cells were cultured in Lonza’s EGM (CC-3162), and pericytes were grown in Immortalized Pericyte Medium (Creative Bioarray, CM-C12030Z).

### CAD Disease-Relevant Stimulation

Primary fibroblasts and primary SMCs were seeded at 100K, primary pericytes were seeded at 80K, and endothelial cells were seeded at 40K per well in 6-well plates. 48h after seeding, cells were treated with growth media containing disease relevant stimulation, with two replicates per condition for each of the following stimulations: low serum (5% FBS in otherwise standard media), oxLDL (100ug/mL in 5% FBS low-serum media), IL-1α (10ng/uL), IL-6 (20ng/uL) + TNFα (10ng/uL), and high glucose (25 mM concentration). On day 3, stimulation media was refreshed and cells were incubated under standard conditions for an additional 1 prior to harvesting for Mint-ChIP ^33^ or ATAC-Seq ^88^.

### Generation of Constitutively Expressing dCas9-KRAB CRISPRi Cell Lines

We generated constitutively expressing dCas9-KRAB cell lines – using immortalized human vascular fibroblast (ABM, T0446) and human coronary artery smooth muscle cells (Applied Biological Materials Inc., T0557) – as previously described ^47^. Briefly, cells were spinfected (at 930 x g for 150 minutes) with lentivirus carrying dCas9-KRAB construct in the presence of polybrene (10ug/mL). 24 hours after transfection, the media was replaced to allow cell recovery and to get rid of polybrene. 48 hours after transfection, selection was done by treating cells with hygromycin (50 ug/mL) for 5-8 days until all corresponding negative control cells were eliminated. Cells were then pooled and expanded for freezing, and dCas9-KRAB activity was tested after the first passage using methods described in the section “CRISPRi activity assay.”

### CRISPRi Activity Assay

Before spinfection for our unbiased *9p21.3* CRISPRi screen, CRISPRi activity of dCas9-KRAB cell lines was tested using Cellecta’s CRISPRiTest dCas9-Repressor assay kit. Cells were transfected according to manufacturer instructions without centrifugation. After 48 hours. fluorescence was measured on a CytoFLEX S. The ratio of GFP/RFP in dCas9-KRAB cells relative to the parental line was calculated using FlowJo software and used to assess effective repression.

### Lentivirus Preparation

We generated Lentiviral cells by transfecting HEK293T cells at about 60% confluency with 1,250 ng sgRNA, and second generation lentiviral packaging vectors 1250 ng psPAX2 (Addgene #12259) and 250 ng of VSVG (Addgene#12259) using TransIT-LT1 transfection reagent. 24 hours later, media was collected. We collected supernatant from these cells after 72 hours, passed it through a 0.45uM filter; the filtrate was used for downstream analysis. Lentiviral transfection for our CRISPRi-MAC-seq screen was performed as described above, using a *9p21.3* densely tiled library (see **Figure 2C**) made by the Genomic Perturbation Platform (GPP) at the Broad Institute of MIT and Harvard.

### Epigenetic profiling and Chromatin States Annotation by ChromHMM

We leveraged the ChromHMM method (version 1.22) for joint-modeling of chromatin states across the vascular cell types (**Figure 1B**). The histone mark signals (H3K4me1, H3K27ac, H3K4me3, H3K27me3) and the ATAC data under the basal condition were used for annotating chromatin states. The preprocessing of the histone mark and ATAC signals included aligning to GRCh38 reference genome using Bowtie2^89^, calling peaks using MACS3^90^ tools, and subsequent removal of peaks with low-confidence. Low-confidence peaks are characterized by any of the following criteria: (i) the fold-change at the peak summit is below the median of all fold-changes; (ii) the −10log_10_P-value (display score) is lower than the 0.75 quantile of all peaks; or (iii) the P-value at the peak summit exceeds 0.05. A joint model was subsequently trained on binarized peaks, utilizing a 200bp bin size. Chromatin states were functionally annotated through manual examination of the ChromHMM annotated states.

### Identification of Novel Vascular Cell Enhancers

For this analysis, we considered enhancers, strong enhancers, and canonical enhancers from our Chromatin-state joint-modelling as Enhancers. Next, BEDTools^91^ was used to merge the enhancer regions, and to intersect these regions with enhancer regions from the Roadmap project (annotated as genic enhancer1/2, active enhancer 1/2, or weak enhancer) ^31^. These resulted in identifying enhancer regions unique to vascular cells and those previously reported in the Roadmap project.

### Super Enhancer Calling by Rank Ordering of Super Enhancers (ROSE)

The ROSE pipeline ^40,92^ was utilized to rank enhancer regions and identify potential super-enhancer regions. In this process, the enhancer annotations from ChromHMM joint-modeling were used to identify and extract enhancer regions from the basal H3K27ac signals from our vascular cell wall samples (**Figure 1B**). Next, the ROSE pipeline was used to score these regions and to define a cutoff for super-enhancer annotation. Every region ranked above the cutoff is considered as a super enhancer. The pipeline used H3K27ac signals from a whole-cell extract as the control.

### Annotation of Enhancers Within and Around *9p21.3* Haplotype

To annotate enhancer within the *9p21.3* CAD haplotype, we intersected ChromHMM ^37^ annotated enhancers along with ATAC-seq and H3K27ac ChIP-Seq data with with 50 coronary artery disease (CAD) *9p21.3* SNPs. We identified 12 enhancers spanning the *9p21.3* Haplotype and annotated them as Enhancers (E1 to E12).

### Estimating Genetic Heritability

We prioritized cell types mediating risk for *9p21.3* CAD associated SNPs by LD score regression (LDSC) analysis, as previously described ^93^. Briefly, we integrated GWAS summary statistics for different cardiometabolic traits – including including CAD ^62^, Ischemic stroke ^94^, Total cholesterol ^95^, Triglycerides ^96^, T2D ^63^ and control trait like lefthandedness ^97^ and schizophrenia ^64^ – with RNA-seq data set at basal state and in response to CAD relevant stimulatory conditions (**Figure 1B**). At the basal state, gene sets for each vascular cell type were obtained using t-statistics ^45^ – to calculate a t-statistic for every gene by fitting an Ordinary Least Square (OLS) model and comparing its expression against every other cell. Genes were ranked by the absolute value of their t-statistics and we utilized top 10% expressed genes for heritability analysis (**Figure S2J**). Additionally, for cells exposed to CAD-relevant stimulatory conditions, we utilized differentially expressed genes (DEGs) relative to the control treatment to perform heritability enrichment analysis for the CAD relevant stimulatory condition groups. For this analysis, we considered -log_10_(p-adj) ≥ 3.35 as significant.

### Densely Tiled *9p21.3* Guide Library

Guides targeting enhancer regions and variants were designed using CRISPick^98,99^ For our densely tiled *9p21.3* library, 3-5 guides were generated for each of the 50 CAD risk SNPs within the haplotype, and 15 guides were generated for each of the 12 annotated enhancer regions. 36 locus-binding negative control guides, targeting a non-genic region negative for enhancer marks (in vascular cells profiled) near the *9p21.3* locus between coordinates Chr9:22174501-22175087 (hg38), were included in the library. 13 non-targeting guides, which don’t match any specific genomic position, were also included (**Table S6**). Lentiviral sgRNAs were cloned and generated by the Broad Genetic Perturbation Platform (GPP), and stored in an arrayed format in 96-well plates, with targets randomized across plates. All 13 non-targeting guides were included on every plate.

### MAC-Seq (Multiplex Assay for Cells Sequencing)

MAC-Seq was performed as previously described ^46^, with slight modifications: (i) Due to the 96-well format, only the first 96 MAC-Seq barcodes were used (lysis volumes were unchanged); (ii) SPRIselect beads (Beckman Coulter) were used in place of AMPureXP beads at every step; (iii) sonication was performed using a Covaris LE220Rsc, with the settings 200W peak incident power, 25% duty factor, 50 cycles per burst, and 85 second duration. Library QC and quantification including Tapestation, Qubit, and KAPPA PCR was performed as described ^46^ and before sequencing on a NextSeq2000 using NextSeq P3-100 flow cells. Accordingly, 750pM of the prepared library was used for sequencing, omitting PhiX spike-in, and the custom read 1 primer was diluted to 0.3uM in Buffer HT1 before use.

### Analysis of MAC-seq: BCL to Fastq, Barcode Demultiplexing, and QC

After sequencing, the raw base-call (BCL) were demultiplexed using bcl2fastq (version 2.20) to generate FASTQ files. FASTQs were then aligned to the reference human genome (GRCh38) using STARsolo (STAR version 2.7.10a) by specifying STAR --soloType CB_UMI_Simple --soloCBwhitelist [/path/to/barcodes.txt] --soloCBstart 1 --soloCBlen 10 --soloUMIstart 11 --soloUMIlen 16 --quantMode GeneCounts --soloFeatures Gene --readFilesCommand zcat. The reads were demultiplexed based on the MAC-seq well barcodes (**Table S26**) to generate the raw UMI count matrices. Mitochondrial and ribosomal gene fraction were calculated per well, gene counts were TPM-normalized, and expression of *9p21* genes in biological replicate plates were compared (**Figure S10**).

### Analysis of MAC-seq: Batch Correction and Permutation Analysis

To correct for batch effect in gene expression levels across plates, expression of *9p21.3* genes were z-score normalized on a per-plate basis. A permutation-based sliding window approach was used to identify strongly repressive regions while accounting for the highly variable number of guides in each window. First, the locus was tiled into 1000-bp windows, staggered every 500bp for 50% overlap, For each *9p21.3* gene, the individual z-score of all guides within each window were averaged to calculate a window’s mean normalized gene expression. Then, the labels of all guides were randomized, a permuted mean expression value was calculated, and this process was repeated for 500 total iterations. To assign significance to observed data relative to the permutation background, a permutation z-score was calculated. For each sliding window, the mean and standard deviation of the 500 permutations were calculated. Then, the mean experimentally observed gene expression was compared with the permutation values to generate a final permutation z-score for that window. A significance level of z-score =-1.96 was used to identify significantly repressed regions.

### Combined Haplotype Test

Combined haplotype tests were performed on RNA profiles using the WASP analysis pipeline^52^. This pipeline was used to evaluate the effects of variants on exon regions of the genes within the *9p21* locus. The results of the pipeline were then aggregated for each gene, encompassing all variants and exon regions. To determine the significance level of the analysis, a binomial test was conducted. This test compared the number of tested conditions against the number of significant tests (P-value< 0.05).

### Assessing the Implication of MTAP in Population-Scale Data

#### FinnGen

We investigated the Olink proteomics data (2025) from FinnGen. The population contains N=1,829 participants from which 1,624 were part of the cardiovascular disease cohort (FinnGen code: FG_CVD [https://r13.finngen.fi/pheno/FG_CVD]). The overlapping sub-population contains 539 cases (50% female, median age 51.9 years), and 1,085 controls (52% female, median age 51.7 years). MTAP Olink expression was then adjusted for age and sex. For cases, age was defined as the age at the event, while for controls, it was the age at the end of follow-up. Wilcox test was used to quantify the differences between the adjusted MTAP expression of the cases vs controls.

#### STARNET

The Stockholm-Tartu Atherosclerosis Reverse Networks Engineering Task (STARNET) ^66^ investigates coronary artery disease (CAD) in seven distinct tissues. The STARNET portal (http://starnet.mssm.edu/) provides differential expression data for target genes, comparing CAD patients (N=600) with a control group (N=250). We utilized the portal to query the genes within the *9p21.3* locus; the log2 fold changes between the disease and control cases and their corresponding p-values were downloaded from the portal.

#### Human Dilated and Hypertrophic Cardiomyopathy

For this study, we utilized data from a previous single-nucleus RNA sequencing study of left ventricle samples ^65^. These samples included 11 hearts with dilated cardiomyopathy (36.36% female, median age 56 years), 15 hearts with hypertrophic cardiomyopathy (33.33% female, median age 51 years), and 16 non-failing hearts (62.5% female, median age 58 years).

We assessed the expression of genes within the *9p21.3* locus. Gene expressions were normalized via log-transformation, and age and sex were regressed out. The residuals from this adjustment were then used for a non-parametric comparison of means. To compare adjusted expressions between patients and control participants, we employed a Wilcoxon test. This analysis was performed on individual annotated cell types and repeated after pseudo-bulking the data. The statistical output of these comparisons is presented in Table S27.

#### Gene Prioritization Using MAGMA

The candidate genes at the *9p21.3* locus were prioritized by using genome-wide association study (GWAS) summary statistics for coronary artery disease (CAD), type 2 diabetes (T2D), and schizophrenia (SCZ). Gene-based association analyses were performed with MAGMA v1.10 (Linux, GCC 5.2) that includes SNP annotation and gene-based analysis steps. SNP locations were annotated to NCBI gene coordinates using 1000 Genomes phase 3 reference pane (hg19 build), and gene-based statistics were computed using GWAS summary statistics with P-values and sample sizes. Gene-level Z-scores and P-values were obtained for each trait, and percentile ranks were calculated for both Z-score and P-values.

#### Genotyping

Vascular cell wall samples were sequenced using the Illumina Global Diversity Array (GDA), yielding 1,904,599 variants per sample. These variants showed a high correlation with the reference genome (r2 = 0.879). Subsequently, the data underwent quality control, phasing (Eagle v2.4), and imputation using the TOPMed Imputation Server, based on the TOPMed r3 reference panel. After imputation, quality control was performed. Variants with a minor allele frequency (MAF) of 0.001 or less and a Hardy-Weinberg equilibrium P-value of 1×10^-20^- or less were removed using the PLINK tool set. Finally, the BCFtools ^100^ was used to remove variants with R^2^ > 0.5, resulting in 7,854,254 variants for downstream analysis.

#### LipocyteProfiler, Fibroblast Activation Experimental Procedure, Imaging, and Analysis

For high-content imaging, primary smooth muscle cells and fibroblasts were seeded in Perkin Elmer (now Revvity) 96-well PhenoPlates at a density of 8,000 cells per well, avoiding wells on the plate perimeter. One day after seeding, cells received either no treatment, DharmaFECT 1 and siRNA control, or DharmaFECT 1 and gene-targeted siRNA (50nM final concentration) for 9p21 gene knockdown (Dharmacon: DharmaFECT 1 Transfection Reagent, ON-TARGETplus Non-targeting Pool #D-001810-10-0010,ON-TARGETplus Human MTAP (4507) siRNA SMARTpool #L-009539-00-0010, ON-TARGETplus Human DMRTA1 (63951) siRNA SMARTpool #L-026122-00-0010,ON-TARGETplus Human CDKN2A (1029) siRNA SMARTpool #L-011007-00-0010, ON-TARGETplus Human CDKN2B (1030) siRNA SMARTpool #L-003245-00-0010, Lincode Human MIR31HG siRNA SMARTpool #R-187931-00-0010, Lincode Human CDKN2B-AS1 siRNA SMARTpool #R-188105-00-0010). Media was changed to standard cell culture media on day two after seeding, and then to Opti-MEM with or without 80ng/mL TGF-β (Sigma Aldrich #T7039) on day three. Media (+/- TGF-β) was refreshed on day four, and on day 5, media was removed and cells were stained using either the LipocyteProfiler or fibroblast activation stain panel. LipocyteProfiler stain was conducted as previously described (CITE LP paper). For the fibroblast activation antibody stain, cells were fixed with 40ul 4% PFA containing 0.5% Triton-X 100 for 20min at room temperature and then stained using the procedure and antibody combination as described ^65^. Plates were imaged using a Perkin Elmer Opera Phenix confocal microscope.

For the fibroblast activation stain, image analysis was performed in Revvity’s Signal Image Artists (SImA) software. Single nuclei were identified by Hoechst signal, and cell bodies were thresholded using a composite COL1a1 and α-tubulin signal. Total cell counts and mean stain intensities per cell were calculated for each channel. αSMA fibers were additionally identified by thresholding the αSMA stain channel for all regions with an intensity value of over 1500. This threshold was able to maximally label in-plane actin fibers while excluding background signal from all wells. The total αSMA fiber area and mean fiber intensity per well were then calculated.

#### Quantification and Statistical Details for Image Analysis

Quantitation was performed by running CellProfiler 4.2.6 on CellProfiler-on-Terra (https://github.com/broadinstitute/cellprofiler-on-Terra). Before segmentation, illumination correction was applied across each image channel (DNA, BODIPYRNA, AGP, Mito) using correction images generated per plate to compensate for uneven illumination. Nuclei were identified using the HOECHST stain, and the watershed method was used to identify whole cells using the Phalloidin/WGA and BODIPY/SYTO14 stains. Cytoplasm regions were identified by removing the nuclei from the Cell segmentations. For each nucleus, cell, & cytoplasm, measurements were collected representing location, size, shape, t5 objects. After feature extraction, the data was filtered by applying automated and manual quality control steps. One well with a total cell count of less than 9000 was removed in CFB as it was a low outlier in its cell type group. Second, every well was visually assessed, and treatments with experimental artifacts were removed as a QC step (Table S28). Next, only siMTAP knockdown wells and non-targeting controls from the same plate were retained. Measurement categories known to be noisy, namely Manders, RWC, and Costes features ^101^, were removed. Additionally, features with zero standard deviation across all images, bounding box metrics, and X and Y coordinate features were removed.

To reduce feature redundancy, we calculated the Pearson correlation (R²) for all feature pairs, creating a distance matrix with 1-R². We then applied complete-linkage hierarchical clustering to this matrix, grouping features into 850 clusters from 3,252, each maintaining a minimum intracluster R² of 0.9. The median of features within each cluster served as its representative signal of the cluster.

Signals were normalized using the inverse normal transformation (R’s RankNorm ^102^) for downstream analysis. A similarity matrix was then generated (using Morpheus ^103^) between all non-targeting control and siMTAP-knockdown wells. For each cell type, two pairwise comparisons were conducted: basal siMTAP vs. basal nontargeting control, and TGF-β siMTAP vs. TGF-β nontargeting control. Statistical significance for each normalized feature was determined using the Wilcoxon rank sum test (Mann-Whitney U), and raw p-values were adjusted to q-values using the q-value package (https://bioconductor.org/packages/release/bioc/html/qvalue.html).

#### siRNA Treatment and RNA-Seq

Parental fibroblasts and smooth muscle cells were grown in twelve-well plates for RNA-Seq. On day one after seeding, cells were treated with siRNA as described above, with 5uL Dharmafect per mL of media and a 50nM final siRNA concentration. Three biological replicates per siRNA were used. On day 5, cells were frozen, and then total RNA was extracted using the Qiagen RNeasy Plus Mini kit and concentration and quality were measured by NanoDrop2000. For each siRNA condition, the two highest quality replicates were selected for sequencing.

Sequencing data were initially processed by the Broad Genomics Platform (GP), where raw sequencing reads were aligned against the human genome (hg19) reference assembly and stored as CRAM files. Subsequently, CRAM files were converted back into FASTQ files using Picard (version 2.9.0, function: SamToFastq). Paired-end reads were then pseudo-aligned to the human transcriptome derived from the hg38 reference annotation (Gencode v44) using Kallisto (version 0.48.0). The resulting transcript-level estimates were aggregated into gene-level counts using the tximport package in R [version 1.24.0, function tximport(fileList, type = “kallisto”, tx2gene = tx2gene, ignoreAfterBar = T)]. The gene-level count matrix was used for differential gene expression (DEG) analysis using the function DESeq() in DESeq2 (version 1.36.0). DEGs with adjusted P-value< 0.05 and |log2 fold change| >=1 were considered significantly DEGs between groups.

#### Pathway and Network Analysis

We performed gene ontology enrichment analysis for significant differentially expressed genes (P-adj<0.05) using over-representation analysis with the gProfiler tool ^104^. We performed network analyses of the enrichment results to find relevant functional modules by creating a weighted network of these enriched GO terms by establishing edges between terms based on shared genes, using the Jaccard similarity measure that reflects the degree of functional overlap between two biological processes. The network was then clustered using the Leiden algorithm ^105^.

The resulting clusters represent cellular processes overrepresented in statistically significant differentially expressed genes during siMTAP knockdown. Within each pathway in a cluster, we can visualize genes as upregulated (red) or downregulated (blue) based on their annotation to that pathway. The size of each pathway denotes its fold enrichment, indicating how well that pathway is enriched in our differentially expressed genes compared to the background. The network analysis and visualization was implemented using R package igraph (v2.1.4) ^106^. To improve visual clarity, only parent GO nodes were included in the network representation when all parent and corresponding child terms were present in the list.

#### *9p21.3* Fine-Mapping

*9p21.3* Fine-mapped data from FinnGen for the trait “Coronary revascularization (ANGIO or CABG)” (exact criteria as previously described) were accessed from the FinnGen browser (https://r12.finngen.fi/region/I9_REVASC/9:21900177-22300177). Briefly, in-sample dosage LD was calculated using LDStore2 ^107^ and then finemapping was conducted using SuSiE with in-sample dosage LD and GWAS summary statistics as input (https://finngen.gitbook.io/documentation/methods/finemapping).

#### Western Blot Analysis

Western blot analysis was performed as previously described ^108^. Briefly, we probed for Cas9 expression in our constitutively expressing dCas9-KRAB vascular cells using specific antibodies (Diagenode; C15310258). For loading control, we utilized GAPDH antibody (Cell signaling, 5174S). Total protein isolation and blotting was done as following previous protocol in the lab ^108^

#### Relative Gene Expression Analysis by qPCR

For validation of siRNA efficiency, the same RNA used for RNA-Seq as described above was also used for qPCR. cDNA was generated from 500ng RNA using ThermoFisher’s High Capacity cDNA reverse transcription kit, and diluted 1:4 in nuclease-free water prior to qPCR. The qPCR reaction mixture was as follows: 5uL Power SYBR™ Green PCR Master Mix, 0.04uL each 100uM forward and reverse primers, 3.92uL nuclease-free water, and 1uL diluted cDNA.

Additional qPCR analysis was performed on independently cloned subset of guides from the 9p21.3 locus screen, cells were transfected in 96-well plates following the transfection protocol previously described. cDNA was generated from direct lysate using Invitrogen Power SYBR Cells-to-Ct kit according to manufacturer protocol.

All qPCR was performed using a BioRad CFX Opus 384 machine, with a denaturation step of 95 °C for 3 minutes and 40 cycles of 95°C for 10 seconds and 60°C for 30 seconds, followed by a melting curve. Relative gene expression was calculated by the 2^−ΔΔCt^ method ^106,109^. Ct values were normalized to TBP expression. Primers used were designed using Primer-BLAST and can be found in Table S29.

#### Human Left Anterior Descending (LAD) Coronary Artery Tissues Collection

Approval for this research was obtained from the Ethics Committees of Zhongnan Hospital of Wuhan University. The study was carried out in accordance with the principles outlined in the Declaration of Helsinki.

Freshly explanted hearts were obtained from orthotopic heart transplantation recipients at Zhongnan Hospital of Wuhan University under protocols approved by the Ethics Committees of Zhongnan Hospital of Wuhan University, with written, informed consent from all participants. Hearts were arrested in cardioplegic solution and rapidly transported on ice from the operating room to an adjacent laboratory. The proximal 5–6 cm segment of left anterior descending coronary artery was dissected on ice, trimmed of surrounding adipose tissue (and in some samples, the adventitia), rinsed in cold phosphate-buffered saline (PBS), and snap-frozen in liquid nitrogen. Frozen tissues were transferred to a single cell library preparation and sequencing laboratory under a material transfer agreement and IRB-approved protocols. All samples were stored in liquid nitrogen prior to further processing.

#### Sample Preparation

All specimens were snap frozen with liquid nitrogen immediately after sampling and stored therein until processing. On the day of the experiment, the frozen tissue was minced into small fragments and transferred to a Dounce tissue grinder. 1 ml of chilled 0.1x lysis buffer (10 mM Tris-HCl pH 7.4 (Sigma-Aldrich, catalog no. T2194), 10 mM NaCl (Sigma-Aldrich, catalog no. 59222C), 3 mM MgCl2 (Sigma-Aldrich, catalog no. M1028), 0.1% Tween-20 (Thermo Fisher, catalog no. 28320), 0.01%NP40 (Sigma-Aldrich, catalog no. 59222C), 0.001% Digitonin (Thermo Fisher, catalog no. BN2006), 1 mM DTT (Sigma-Aldrich, catalog no. 646563), 1% BSA (Sigma-Aldrich, catalog no. A1933), 1 U/μl RNase inhibitor (Promega, catalog no. N2615)) was added, and the tissue was immediately homogenized 15 times slowly and smoothly with loose (A) pestle. After a 5-minute incubation on ice, the mixture was pipetted 10 times with a wide-bore pipette tip, followed by another 10-minute incubation on ice. Then, 1 ml of chilled Wash Buffer (10 mM Tris-HCl pH 7.4, 10mM NaCl, 3mM MgCl2, 0.1% Tween-20, 1mM DTT, 1% BSA, 1U/μl RNase inhibitor) was added to the lysate. The resulting suspension was filtered through a 70 µm Flowmi cell strainer into a 5 ml tube and centrifuged at 500 g for 5 minutes at 4 °C. The supernatant was carefully removed without disturbing the nuclear pellet, which was then resuspended in chilled PBS-RNase inhibitor (1x PBS supplemented with 1 U/µl RNase inhibitor (Sigma, 3335399001)). If cell debris and large clumps were observed, pass through a 40 µm cell strainer again.

Nuclei were then fixed by adding formaldehyde (Sigma-Aldrich, catalog no. F8775) to a final concentration of 1.5%, followed by a 10-minute incubation at room temperature. The fixed nuclei were pelleted by centrifugation at 600 g for 5 minutes at 4 °C. The supernatant was discarded, and the pellet was washed twice with 1 ml of PBS-BSA-RNase inhibitor, with centrifugation at 500 g for 5 minutes at 4 °C after each wash. After the final wash, the supernatant was removed, and the pellet was resuspended in a chilled diluted nuclei buffer (PN-2000153, 10x Genomics) and kept on ice.

### Single-Cell RNA Sequencing

#### Single cell Multiome (3’RNA&ATAC-seq) libraries were processed using the UDA-seq method

For each channel, 300,000 nuclei were loaded and subjected to 96-well post-indexing. Briefly, nuclei were subjected to tagmentation reaction, partitioned and barcoded using the 10x Genomics Chromium Controller with the Chromium Next GEM Single Cell Multiome ATAC + Gene Expression (10x Genomics; PN-1000283), following the manufacturer’s protocol. Immediately after the reverse transcription (RT) reaction, 125 µl of Recovery Agent was added to the GEMs to release the nuclei. The Recovery Agent/Partitioning Oil (pink) was carefully removed from the bottom of the tube. Next, 120 µl of diluted nuclei buffer was added to the remaining 80 µl aqueous phase, mixed thoroughly, and evenly dispensed into a 96-well plate, with each well receiving 2 µl. After brief centrifugation, the plate was incubated in a thermomixer under the following conditions: 85 °C for 5 minutes, followed by a hold at 4 °C. The products were purified using the Dynabeads SILANE genomic DNA Kit (Invitrogen, catalog no. 37012D) according to the manufacturer’s instructions, and eluted in a final volume of 10 µl of EB buffer per well.

Next, well-specific index PCR was performed. A 20 µl of index PCR mix (15 µl NEBNext Ultra II Q5 Master Mix (NEB catalog no. M0544L), 1 µl of 10 µM Partial TSO/IS, 1 µl of 10 µM Truseq-i5 index primer, 1 µl of 10 µM P5 primer, 1 µl of 10 µM Nextare-i7 index primer) was added to each well (note that Truseq-i5 index primer and Nextare-i7 index primer is unique in each well) containing purification products from last step. After brief centrifugation, index PCR reaction was performed as follows: 72 °C for 5min, 98 °C for 3min, and then 7 cycles of [98 °C for 20s, 63 °C for 30s, 72 °C for 1min]; 72 °C for 1min. Half of the indexed PCR products from one 96-well plate were pooled and purified with 1.6x XP beads, then eluted in 160 µl of EB buffer.

##### sc-ATAC-seq Library construction

A 40 µl of the post-indexing product was mixed with 60 µl of ATAC-seq enrichment mix (50 µ1x KAPA HiFi HotStart Ready Mix, 5 µl of 10 µM Partial P5, 5 µl of 10 µM P7 primer). The reaction was amplified as follows: 98 °C for 45s, and then 7-9 cycles of [98 °C for 20s, 67 °C for 30s, 72 °C for 20s]; 72 °C for 1 min. Libraries were size-selected using 0.5-1.4x XP beads and eluted in 25.5 µl EB. Final libraries were sequenced on the MGISeq-T7 platform (MGI Tech Co., Ltd., China) in a 100-bp paired-end mode, with a target depth of 20,000 reads per cell.

##### sn-RNA-seq Library construction

Following cDNA enrichment, 50 ng of cDNA product was used for library construction, following the same procedure as described for UDA-seq. The final libraries were then sequenced on the MGISeq-T7 platform using 150-bp paired-end reads, with a target depth of 50,000 reads per cell.

#### Human LAD Coronary Artery snMultiome Processing and Analysis Pipeline

The preprocessing of snRNA-seq and ATAC-seq data followed a previously described methodology ^110^. Briefly, Read mapping and quantification were performed using Cell Ranger ARC (version 2.0.1). Donor demultiplexing was executed with Demuxlet v2 (https://github.com/statgen/demuxlet) on the alignment files (BAM format) from both snRNA-seq and snATAC-seq data ^111^. This process utilized genotyping information from each donor’s Illumina Infinium Asian Screening Array, which was processed into VCF files imputed either by minimac3 or via the Michigan Imputation Server (https://imputationserver.sph.umich.edu) ^112^. Doublets were identified and removed using Scrublet v0.2.3 ^113^

Data quality control criteria were applied as follows: for the snRNA-seq dataset, we filtered cells with parameters nCount_RNA > 200, nFeature_RNA > 200, and the mitochondrial gene percentage < 10%; for the snATAC-seq dataset, we filtered cells whose TSS enrichment score > 2 and the number of fragments > 1000. Only cells passing the quality control for both snRNA-seq and snATAC-seq were included in subsequent analyses.

We used Seurat v5 (https://github.com/satijalab/seurat) to process the snRNA-seq dataset: (1) Count matrix was normalized using the NormalizeData function (with normalization.method = “LogNormalize” and scale.factor = 10^4^), (2) Top 2,000 highly variable genes were calculated using the FindVariableFeatures function (with selection.method = “vst”), (3) PCA was performed using ScaleData and PCA functions with default parameters. (4) Batch effect was removed using the RunHarmony function (with parameters max_iter = 50 and early_stop = TRUE) implemented in Harmony v1.2.3 (https://github.com/immunogenomics/harmony) ^114^ (5) For visualization, the RunUMAP function was applied to the top 30 PCs ^115^.

We used ArchR v1.0.3 (https://github.com/GreenleafLab/ArchR) to process the snATAC-seq dataset: (1) Tile matrix was generated using the addTileMatrix (tile size 500bp), used by ArchR to quantify chromatin accessibility. (2) The dimensionality reduction was conducted using a latent semantic indexing algorithm, and the top 25,000 variable bins ranked by signal variance were selected as input, (3) Batch effects were removed using addHarmony, (4) UMAP was generated from the 30 dimensions ^116^

To integrate gene expression and chromatin accessibility, we converted the processed ArchR project and Seurat object into a uniform Signac object using ArchR2Signac v1.0.5 (https://github.com/swaruplabUCI/ArchRtoSignac)^117,118^. We then computed a joint neighbour graph that represents both gene expression and chromatin accessibility using the weighted nearest-neighbour method in Seurat. We plotted the joint UMAP embedding with RunUMAP.

To achieve a robust cell type annotation, we applied CHOIR v0.3.0 (https://github.com/corceslab/CHOIR), which employs random forest classifiers in combination with permutation testing throughout a hierarchical clustering tree for the annotation ^119^. This approach enables a statistical identification of clusters corresponding to distinct cell populations. By following the instructions provided by CHOIR, we yielded 20 robust clusters. Three clusters with high expression of fibroblast markers (*DCN*, *LUM*, *COL3A1*, *and LAMA2*) were merged and collectively annotated as fibroblasts, and one cluster was annotated as smooth muscle cells according to markers *ACTA2*, *MYH11*, *MYL9,* and *TAGLN* ^120–123^.

#### Cell Type-Specific Chromatin Accessibility QTL Analysis

Based on the imputed ASA microarray data, we initially grouped cells by their rs1537371 genotypes (CC, CA, and AA) into three categories. Next, we utilized SCENT v1.0.1 (https://github.com/immunogenomics/SCENT) to connect enhancers to target genes ^53^. We used the getMatrixFromProject function (with parameters useMatrix = “TileMatrix” and binarize = TRUE) to build the input cell-by-tile matrix. A raw cell-by-gene expression matrix was used as input. Subsequently, we filtered tiles that were opened or genes expressed in at least 0.1% of cells for each group. Then, we performed SCENT on each *9p21.3* tile-gene pair using the SCENT algorithm (with regr = “poisson” and bin = FALSE). To account for potential confounders, this analysis was adjusted for sample ID, age, mitochondrial gene percentage, log-transformed number of counts, TSS enrichment score, and log-transformed number of fragments.

## Acknowledgements

MC is the Weissman Family MGH Research Scholar Class 2024-2029. This work was supported by the Novo Nordisk Foundation (NNF21SA0072102), UM1DK126185, 2 RCR DK116691, P30 DK040561, and by a sponsored research agreement with Bayer. The image analysis reported in this paper was performed in collaboration with the Center for Open Bioimage Analysis (COBA) which is supported by National Institute of General Medical Sciences NIH P41 GM135019. We want to acknowledge the participants and investigators of the FinnGen study.

## Author Contributions

Conceptualization, T.N.A. and M.C.; Data curation, T.N.A, B.S., Y.B.C., A.W.H, and H.D.; Formal analysis, T.N.A., B.S., Y.H.,Y.B.C., R.K., T.R.J, and H.D.; Funding acquisition, M.C.; Investigation, T.N.A., B.S., M.P.R., A.M.H.,C.M.W., A.V.K., T.R.J.,H.D., and M.C.; Methodology, T.N.A., B.S., Y.H., Y.B.C., M.M., R.K., K.J.K., and H.D.; Resources, T.N.A, M.W., M.P.R., A.V.K., M. Chaffin., K.J.K., H.D., and M.C.; Software, T.R.J. and H.D.; Supervision, T.N.A., C.B.E., T.R.J., H.D., and M.C.; Validation, T.N.A., B.S., and H.D.; Visualization, T.N.A., B.S., Y.B.C., M.M., R.K.,H.D., and M.C.; Writing - Original draft, T.N.A., B.S., Y.H., Y.B.C., M.M., R.K., H.D., and M.C.; Writing - review & editing, T.N.A., B.S., Y.H., Y.B.C., M.M., R.K., S.A., A.W.H., M.W., A.V.K., C.B.E., M. Chaffin., JL, HS, Y.Z., D.C., L.J., Y.L., J.W., H.Z., T.R.J., H.D. and M.C.

---

**Figure S1.**
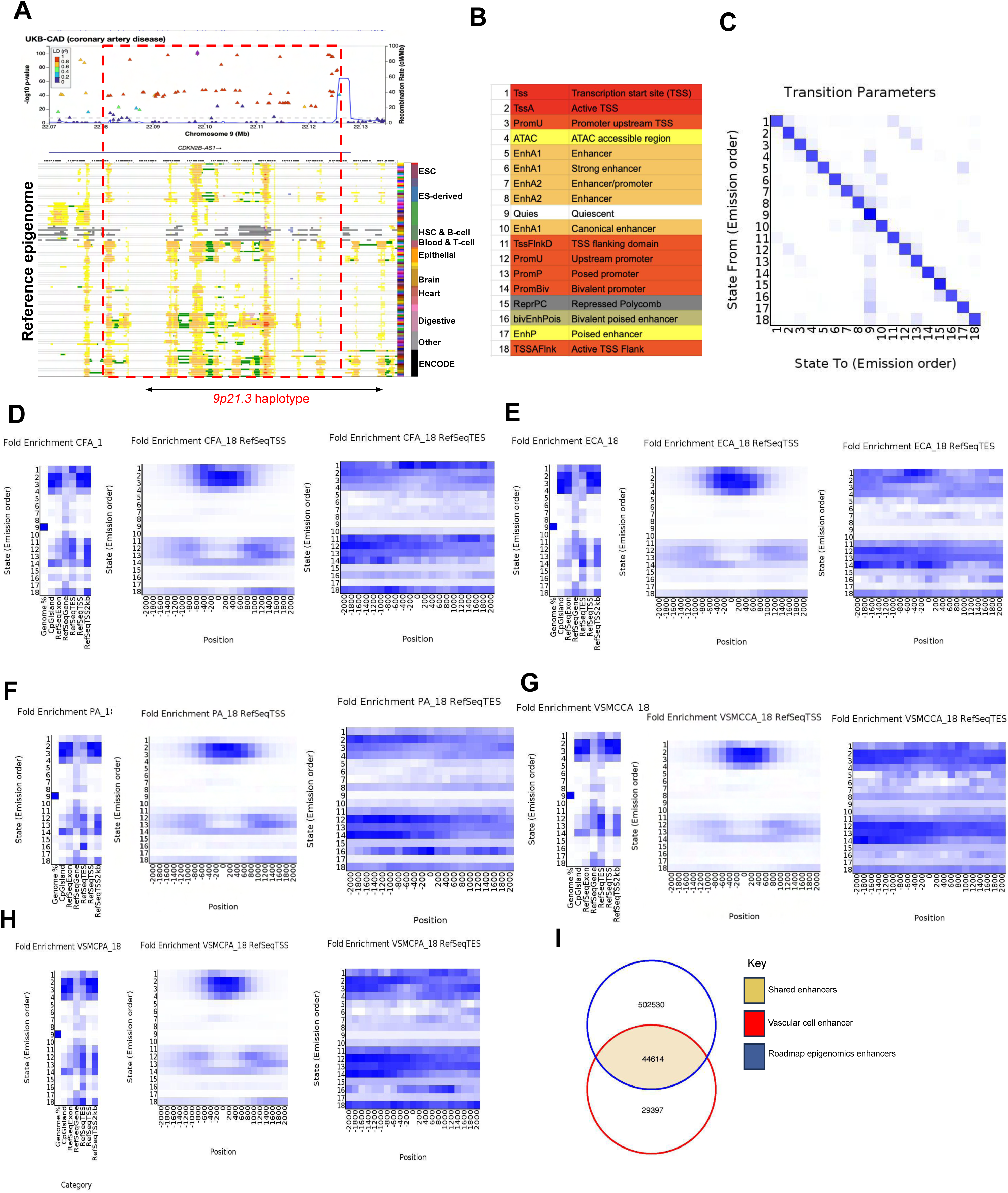
Genome-wide chromatin state annotation of vascular cells by chromHMM (A) Publicly available chromatin state annotation using the ENCODE as well as Roadmap epigenome 111 available data sets. Overlaid above is the UKBB LocusZoom plot for CAD. (B) Candidate state descriptions for each ChromHMM-annotated vascular cell chromatin state, with accompanying state abbreviation. (C-H) Genomic annotations per cell type are displayed in distinct panels. The left heatmap illustrates the fold enrichment overlap for various genomic annotations. A darker blue indicates a higher fold enrichment, relative to a column-specific coloring scale. Next to this, a heatmap displays the fold enrichment for each state within 200-bp bins, spanning 2 kb around RefSeq transcription start sites (TSSs). A darker blue indicates greater fold enrichment, with a single color scale applied across the entire heatmap. The heatmap to the right shows the information but for transcript end sites. (I) Venn diagram showing shared enhancers – with Roadmap epigenome project – and novel enhancers identified by chromHMM annotation of vascular cells

**Figure S2.**
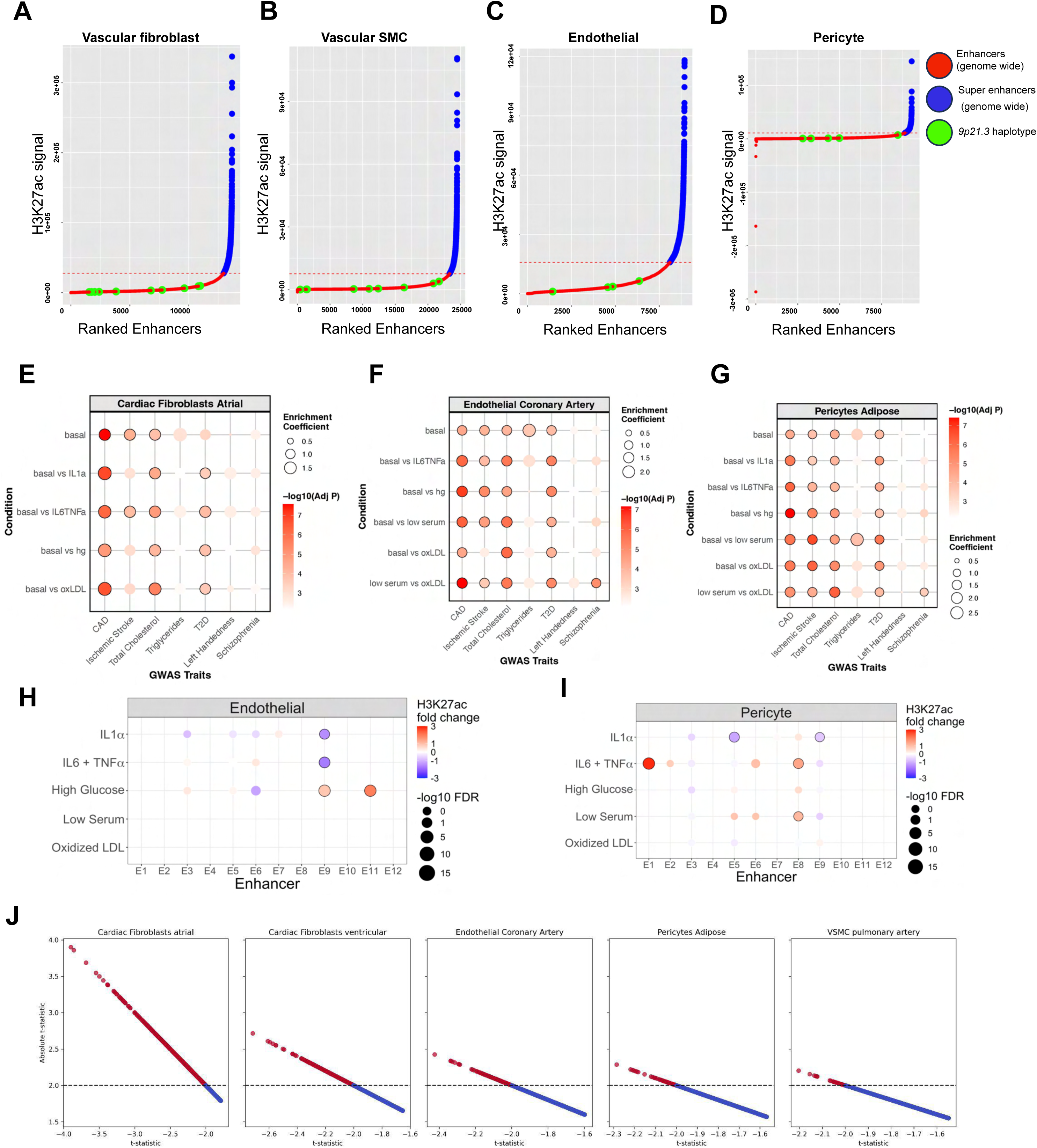
Characterization of *9p21.3* enhancers and identification of cell-types mediating risk for CAD (A-D) Plots showing super enhancers (blue), enhancers (red), and enhancers within *9p21.3* haplotype (green). (E-G) Dot plots showing enrichment for CAD heritability at baseline and in response to CAD-relevant stimulatory conditions, including Inflammatory (IL-1α and IL-6 & TNF-α) atrial derived fibroblast, coronary artery derived endothelial cells, and adipose derived pericytes. Solid dark lines around each dot represent significance at a threshold of -log_10_(adjusted P-value) ≥ 3.35. (H-I) Dot plots showing the impact of CAD-relevant stimulatory conditions on enhancer activities (proxy of H3K27ac) at *9p21.3* locus in endothelial cells and pericytes. Fold changes are indicated as Red (increased fold change) or Blue (Reduced fold change) and solid dark lines around each dot represent significance at a threshold of -log_10_(FDR) ≥ 1.3. (J) T-statistic versus absolute t-statistic (y-axis) for gene expression across five primary human vascular cell types under basal conditions. Differentially expressed genes, representing the top 10% by absolute t-statistic, are highlighted in red; all other genes are in blue. The dashed line indicates a high absolute t-statistic threshold.

**Figure S3.**
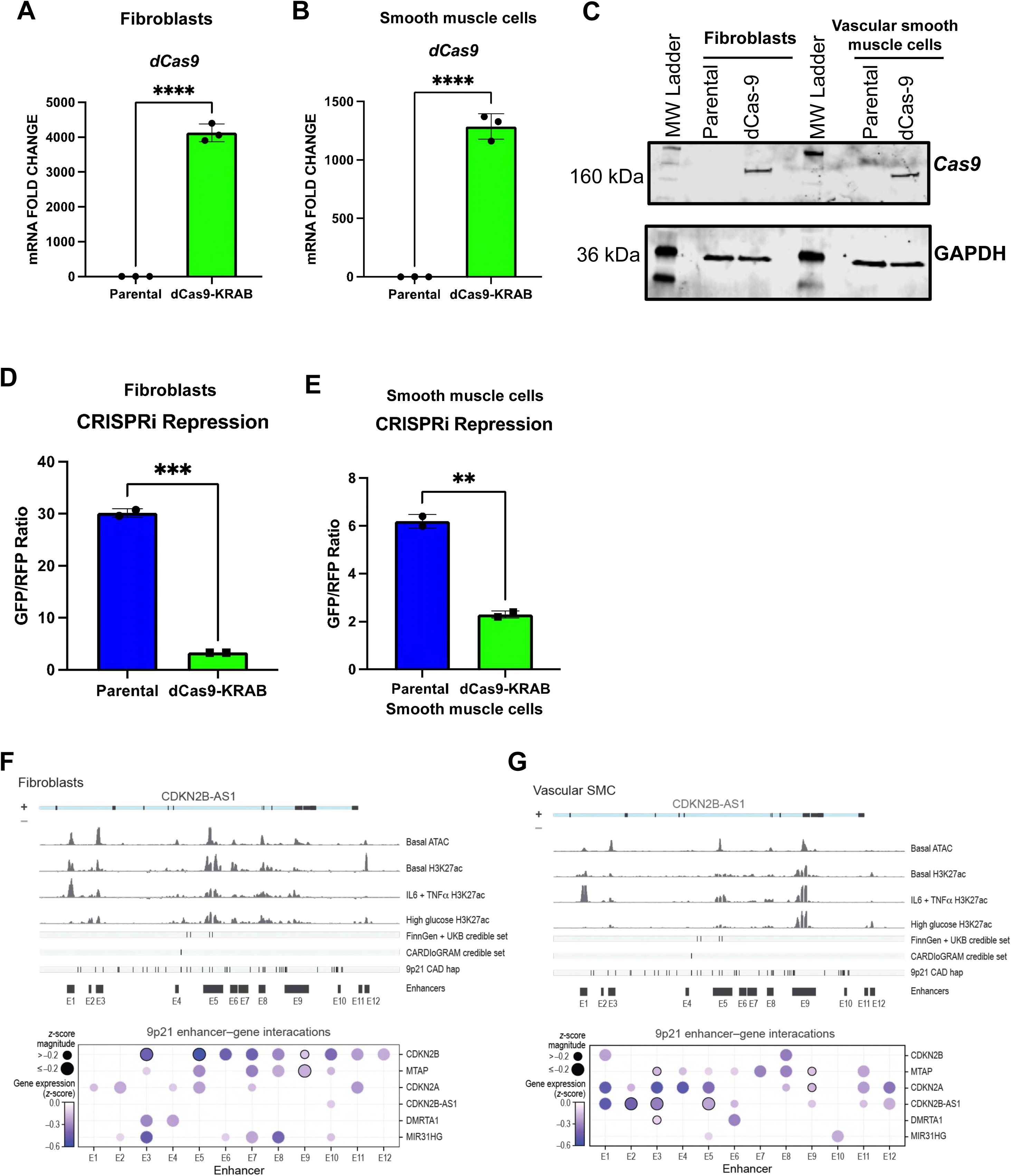
Generation of constitutive expressing dCa9-KRAB (A-B) Bar graph showing RT-qPCR detection of dCas9-KRAB mRNA expression in both control parental cells and the constitutive dCas9-KRAB expressing vascular fibroblast (n=3) and smooth muscle cells (n=3). A t-test was used to identify significant differences between control vascular parental lines and dCas9-KRAB expressing vascular cells. P-value< 0.05 is considered significant. (C) Western blot analysis showing detection of Cas9 protein in the dCas9-KRAB cell lines used in this study. (D-E) Bar graphs showing dCas9-KRAB repression activities. A t-test was used to identify significant differences between control vascular parental lines and dCas9-KRAB expressing vascular cells. P-value<0.05 is considered significant. (F-G) Dot plots showing aggregated effects of *9p21.3* enhancer guides on expression of cis-expressing genes in vascular fibroblast and smooth muscle cells. Dark border circles indicate independent signal validation by qPCR, which was conducted for enhancers E2, E3, E5, and E9. In this plot, we overlaid epigenetic enhancer signatures under basal and disease-relevant stimulatory conditions to show relative position of annotated *9p21.3* enhancers to epigenetic signatures around *9p21.3* haplotype.

**Figure S4.**
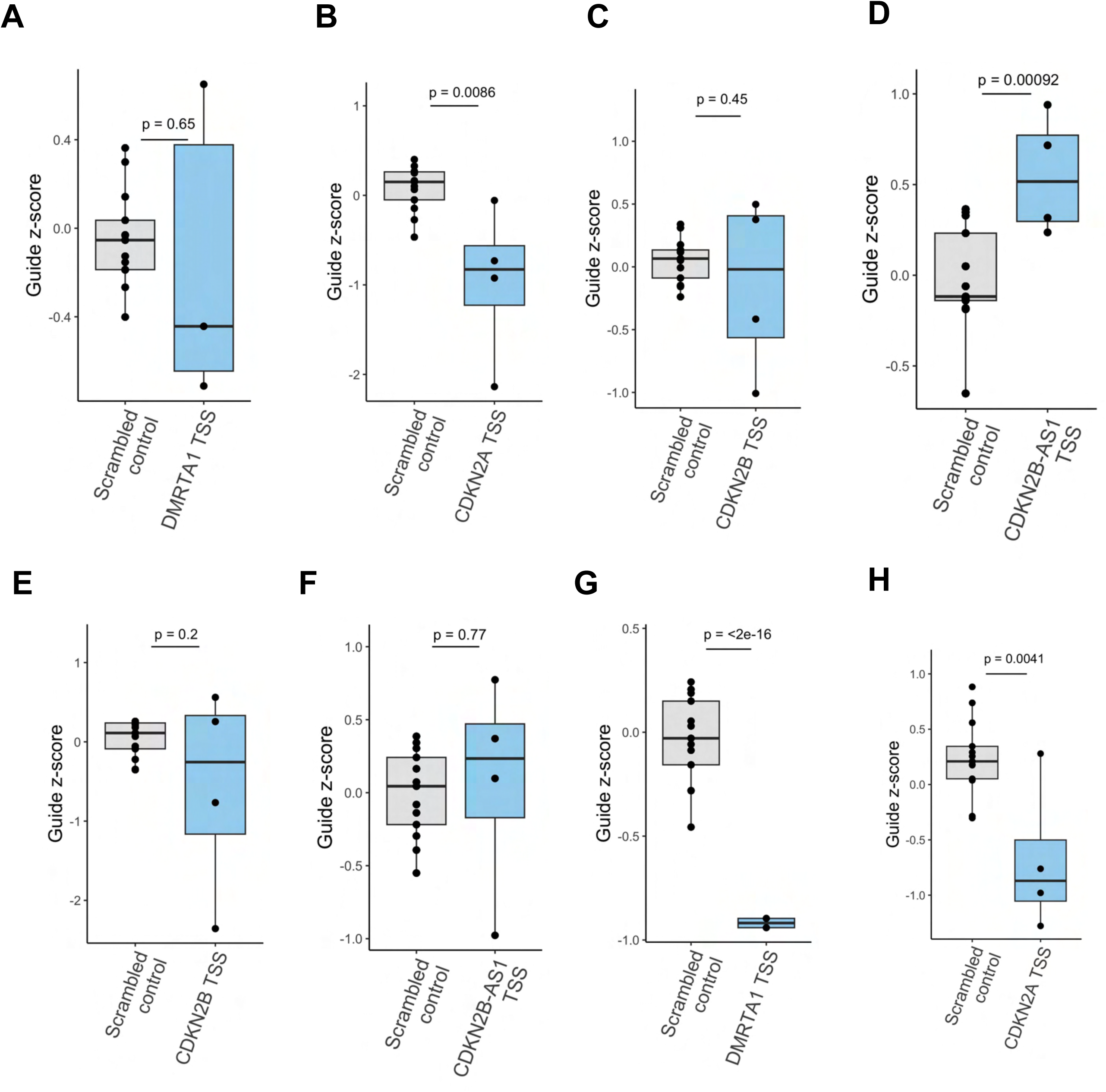
Validation of TSS knockdown of cis-expressing *9p21.3* genes and evaluation of guide efficiency from z-score normalized CRISPRi-MAC-Seq gene expression data. (A-D) Box plot showing knockdown efficiency of *9p21.3* cis-expressing gene vascular fibroblast TSS knockdown efficiency. Analyzed by t-test with a significance threshold of P-value<0.05. (E-H) Box plot showing knockdown efficiency of *9p21.3* cis-expressing gene vascular smooth muscle cell TSS knockdown efficiency. Analyzed by t-test with a significance threshold of P-value<0.05.

**Figure S5.**
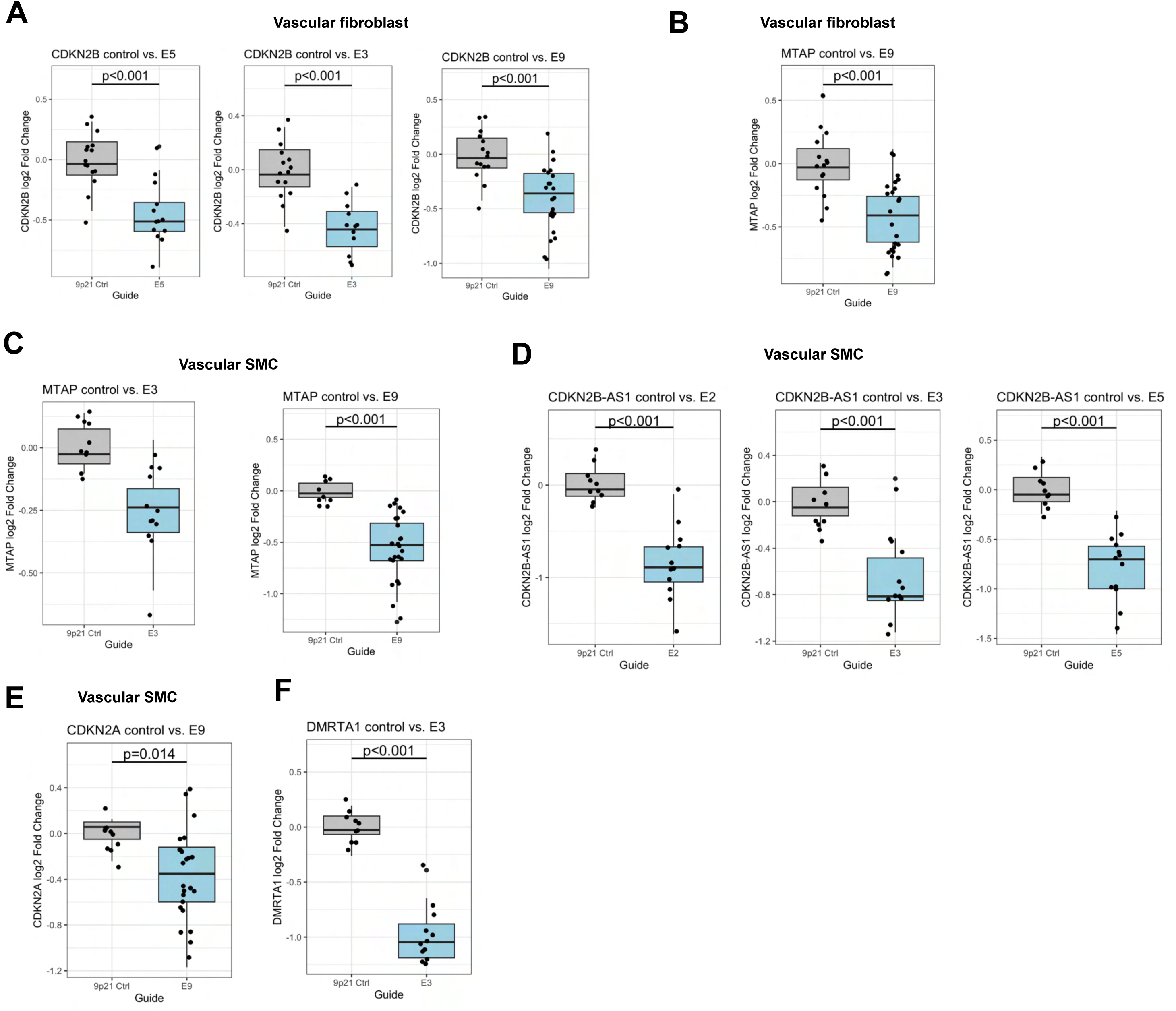
Validation of *9p21.3* enhancer-2-gene (E2G) connection using a direct-lysis qPCR method (Invitrogen two-step Cells-to-Ct). (A-B) Box plot showing effect of 9p21.3 enhancers on cis-gene expression in vascular fibroblasts. Analyzed by t-test with a significance threshold of p<0.05. Guides from the larger screen targeting enhancers E2, E3, E5, and E9 were independently cloned for this validation experiment. (C-F) Box plot showing effect of 9p21.3 enhancers on cis-gene expression in vascular smooth muscle cells. Analyzed by t-test with a significance threshold of p<0.05. Guides from the larger screen targeting enhancers E2, E3, E5, and E9 were independently cloned for this validation experiment.

**Figure S6.**
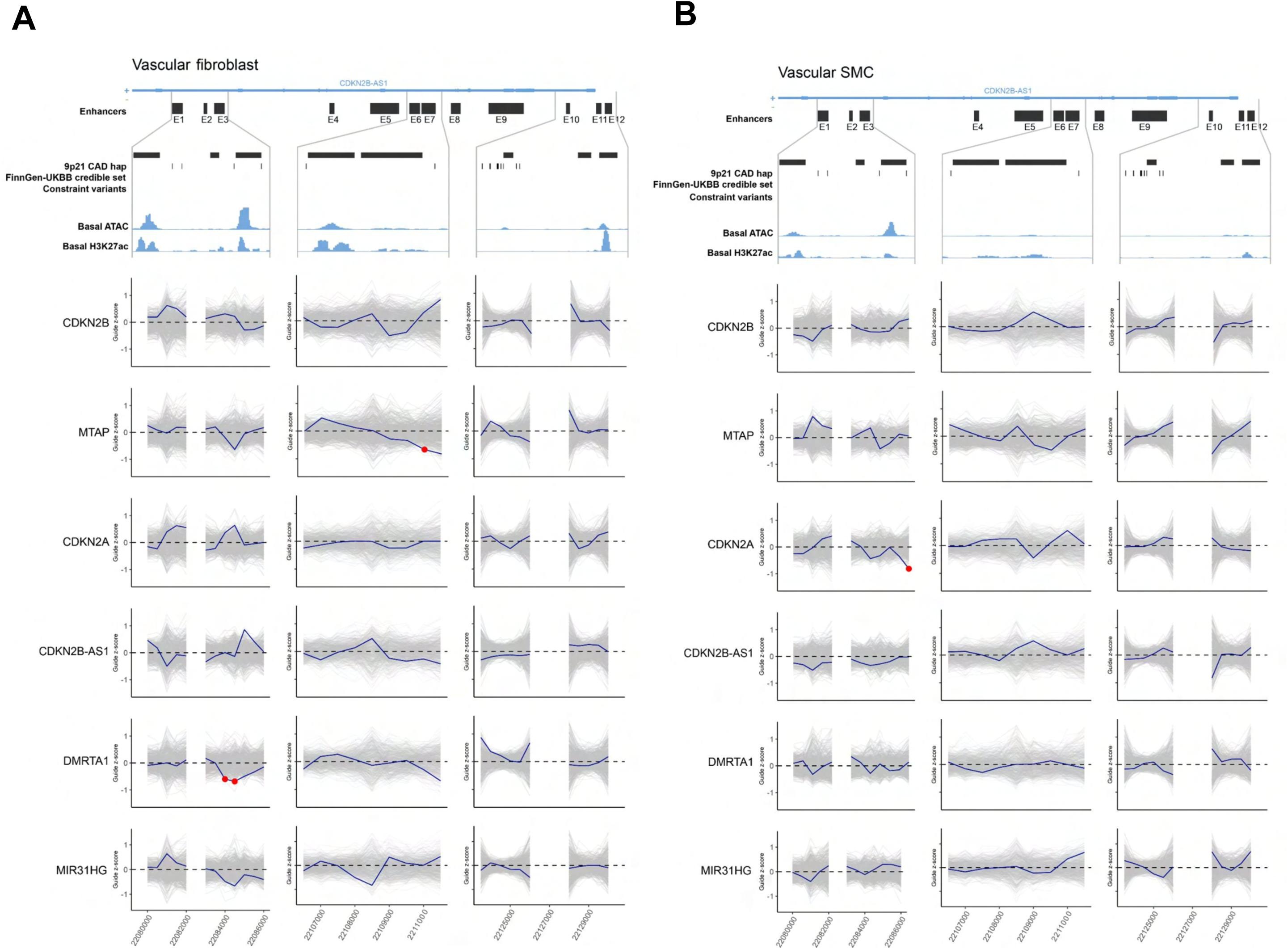
Plot shows a sliding window approach that aggregates sgRNA transcriptional effects across the *9p21.3* annotated enhancers, including E1, E2, E3, E6, E7, E10, E11, and E12 in (A) vascular fibroblast, (B) Vascular smooth muscle cells. To assign significance to observed data relative to the permutation background, a permutation z-score was calculated. A significance level of z-score=-1.96 was used to identify significantly repressed regions – illustrated as a red dot on the plots.

**Figure S7.**
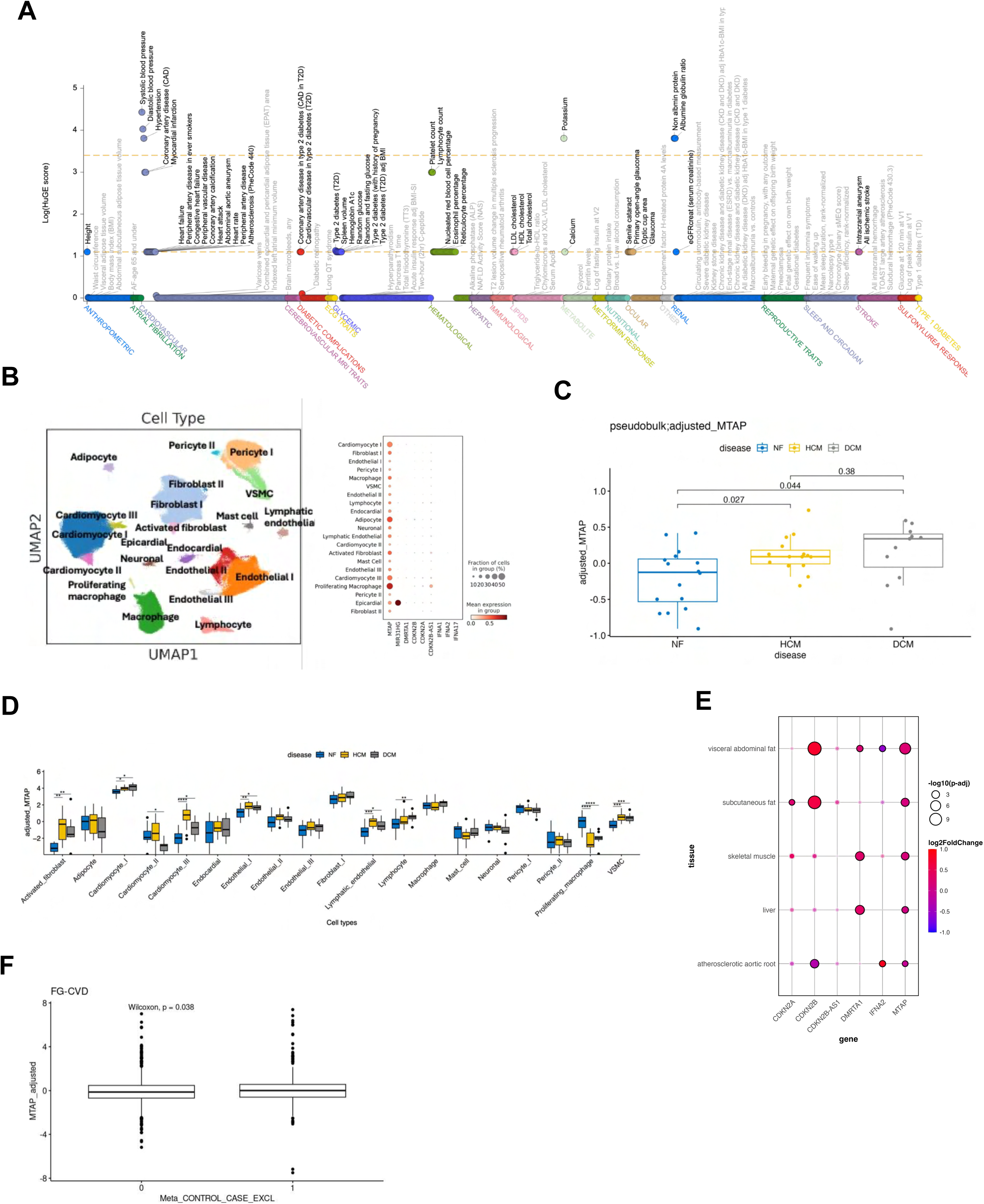
Translational relevance of MTAP in large-scale population (A) left: Uniform manifold approximation and projection (UMAP) of 592,689 single-nucleus RNA profiles from left ventricle samples from 42 individuals ^2^. Right: Expressions of the genes within the 9p21 locus in the 21 identified cell types. The dot plot highlights the expressions of MTAP across these cell types. (B) *MTAP* expression was analyzed in non-failing (NF) participants and compared to patients with hypertrophic cardiomyopathy (HCM) and dilated cardiomyopathy (DCM) using box plots. Significant differences were observed when comparing pseudobulk expressions. Next panels show similar results using the expressions in the fibroblast and VSMC cell types. The trend of increased expression is not significant in fibroblast cells, however, a significant trend was observed in VSMC. (C) Data from STARNET showing tissue-specific changes in 9p21 gene expression in CAD cases vs. control. *MTAP* expression was significantly changed across multiple tissues. (D) Analysis of MTAP expression from the FinnGen Olink proteomics database showing a significant (p=0.038) increase in MTAP expression level in the circulating blood of CAD cases (mean=0.047, 1st quartile= −0.665, 3d quartile=0.520) relative to control (mean= −0.023, 1st quartile= −0.672, 3d quartile= 0.493). (E) To demonstrate the gene prioritization is disease specific, we repeat the analysis using a schizophrenia GWAS. ZSTAT of none of *9p21* genes pass the 0.95th percentile cutoff.

**Figure S8.**
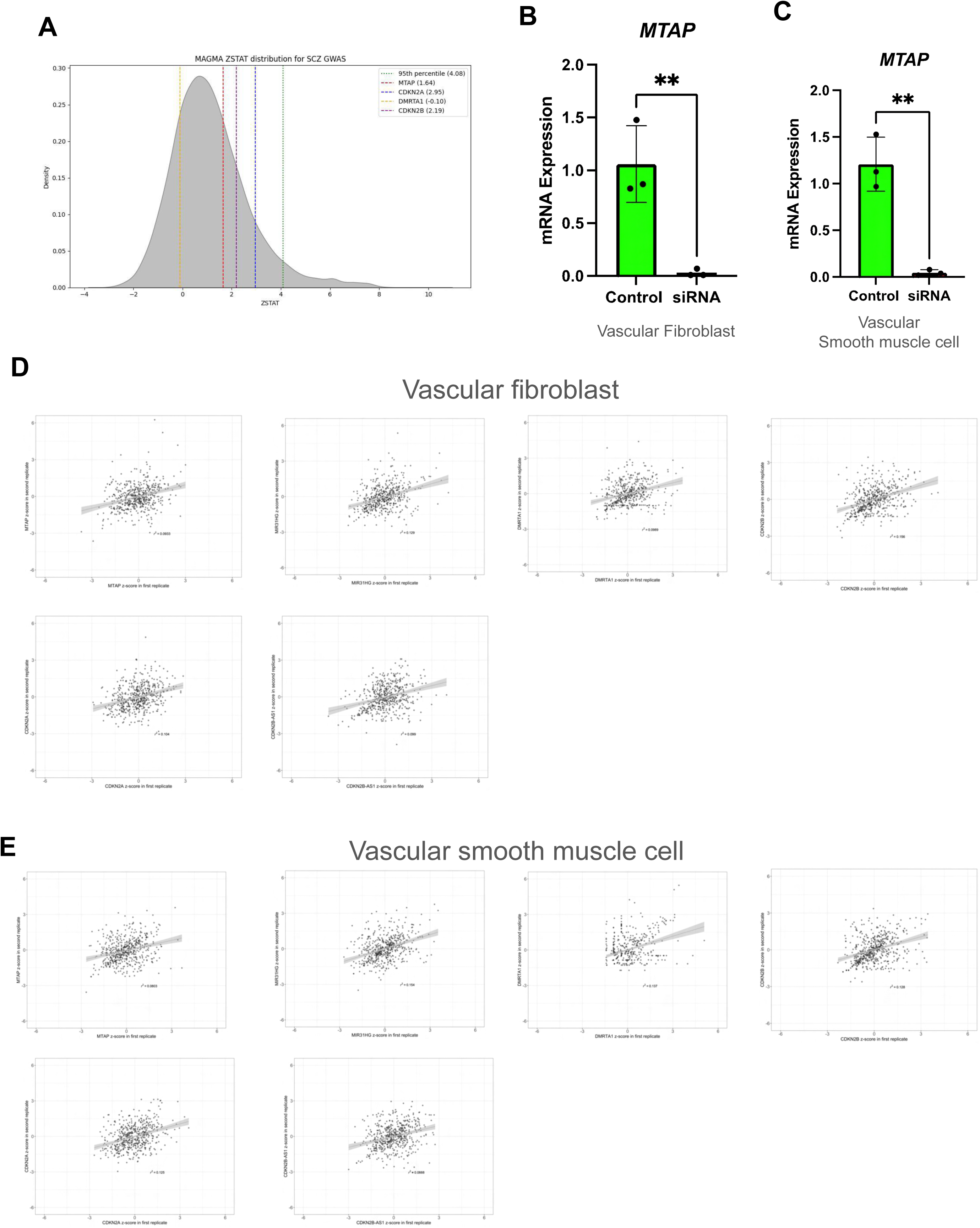
Convergent evidence for MTAP as a driver CAD gene. (A) Density plot from MAGMA showing analysis on Schizophrenia disease trait. (B-C) Validation of siRNA knockdown of *MTAP* in samples sent for bulk RNA-Seq

**Figure S9.**
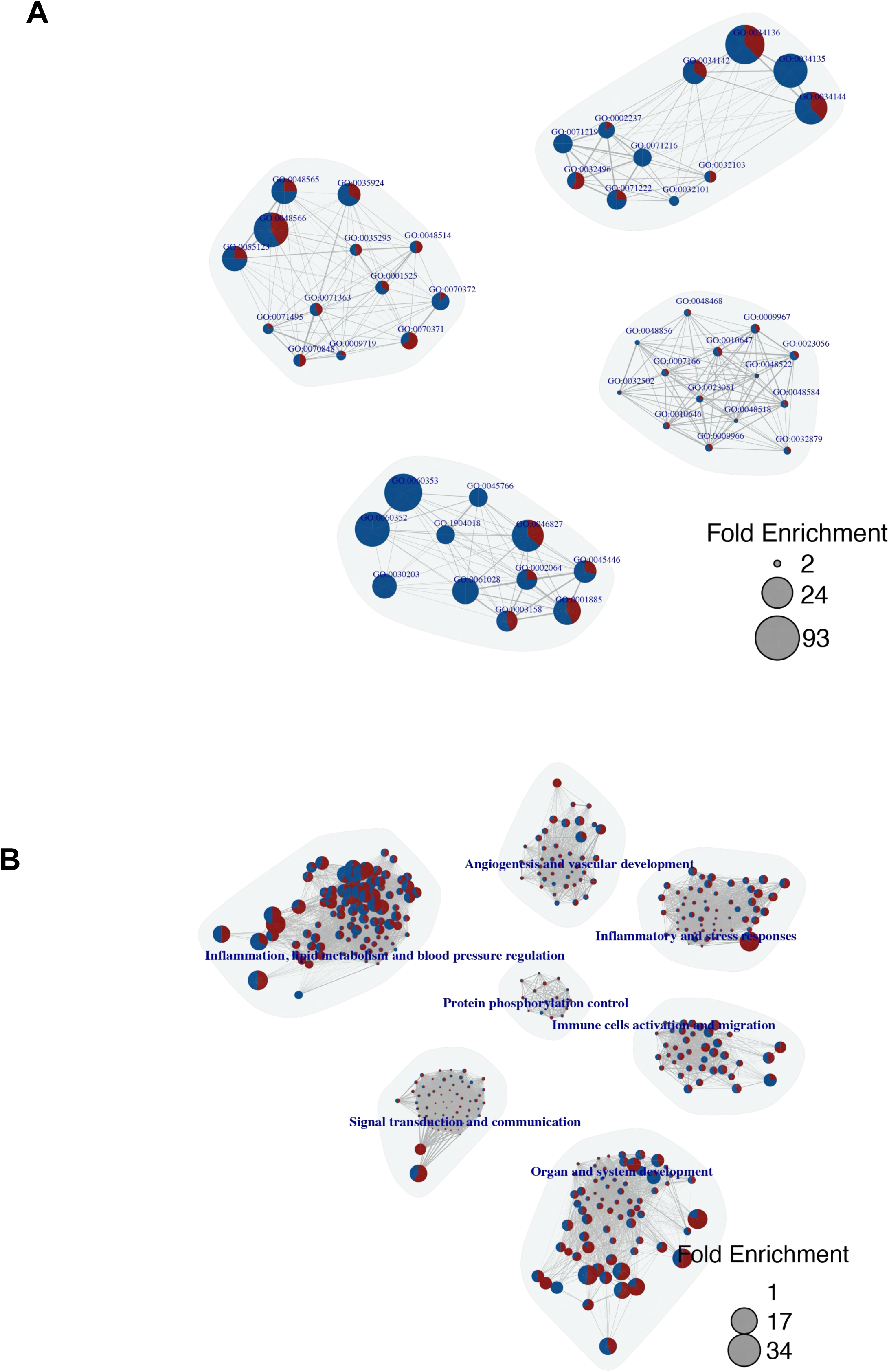
Network analysis of gene ontology pathways impacted by siMTAP knockdown in vascular fibroblast and smooth muscle cells, showing interconnected networks of disease-relevant modules in (A) Vascular smooth muscle cells. (B) Vascular Fibroblasts.

**Figure S10.**
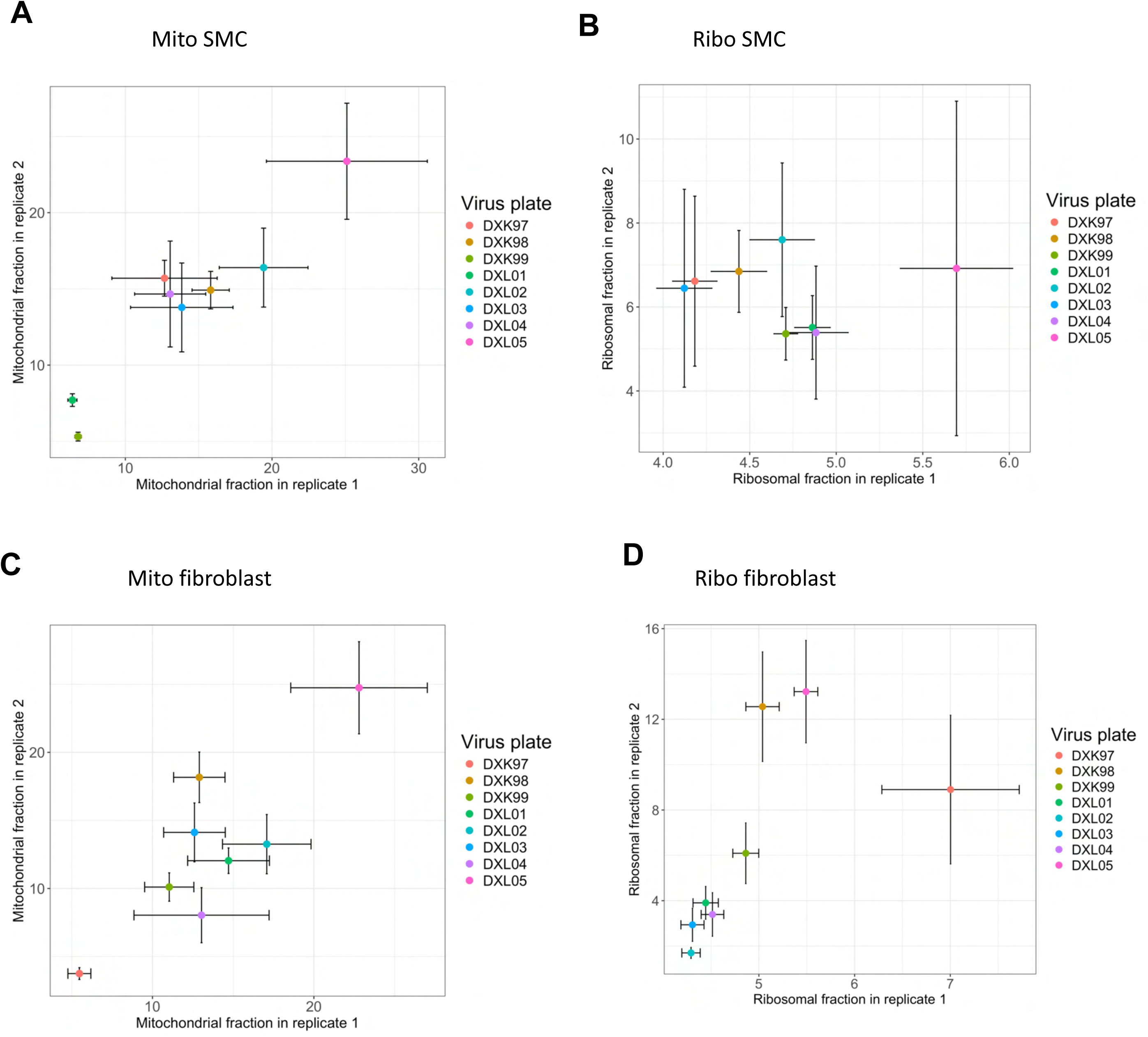
QC showing overall mitochondrial and ribosomal gene fraction for our CRISPRI-MAC-seq screen. For all plots, the x-axis shows the first biological replicate for each of 8 MAC-Seq arrayed plates, and the y-axis shows the second replicate. (A) Mitochondrial fraction (%) by plate with standard deviation in vascular smooth muscle cells (B) Ribosomal fraction (%) by plate with standard deviation in vascular smooth muscle cells. (C) Mitochondrial fraction (%) by plate with standard deviation in vascular fibroblasts. (D) Ribosomal fraction (%) by plate with standard deviation in vascular fibroblasts.

**Figure S11.**
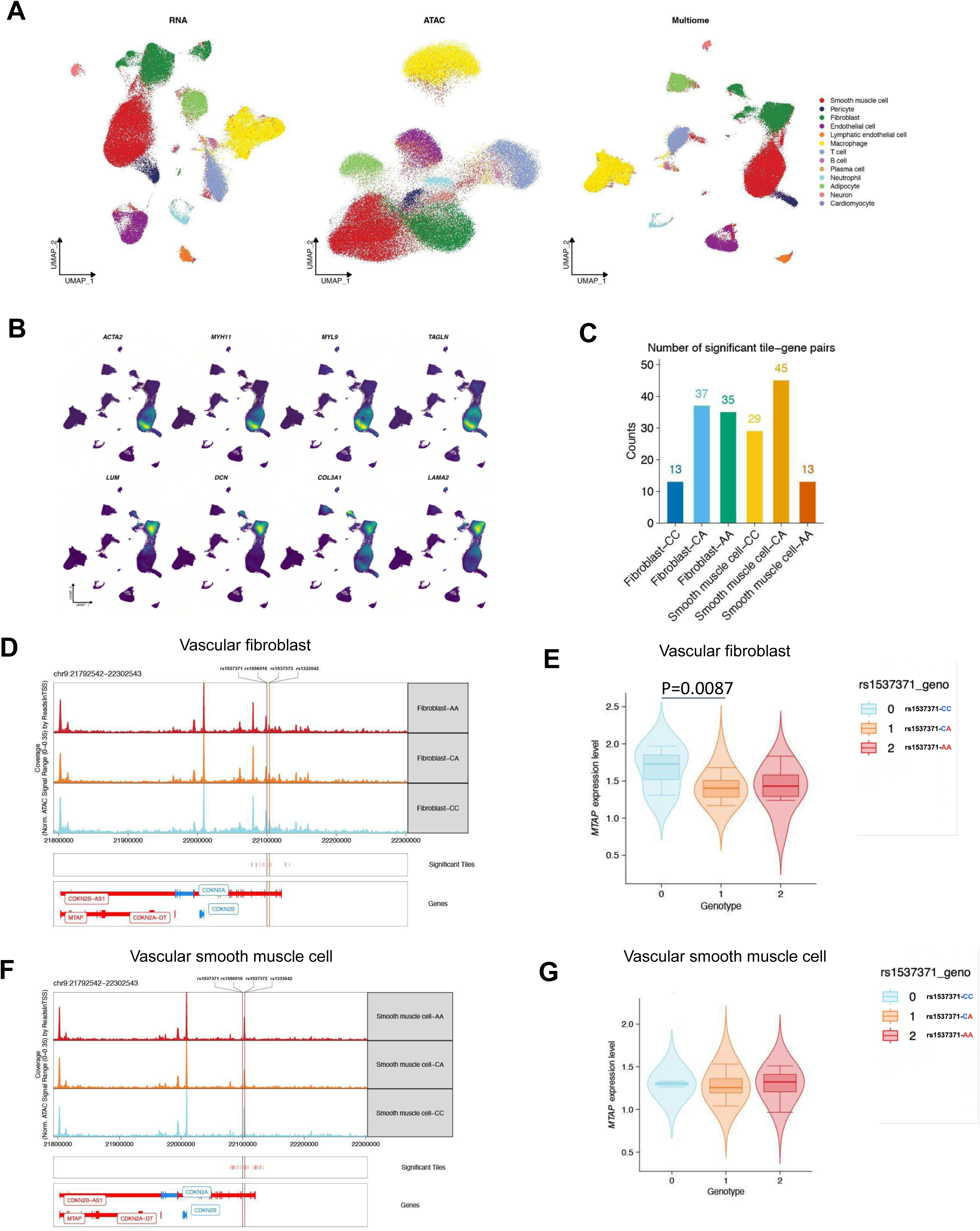
scMultiome analysis of Left anterior descending (LAD) coronary artery. (A) Uniform manifold approximation and projection (UMAP) of scMultiome (scATAC-seq + scRNA-seq) data from Human left anterior descending (LAD) coronary artery, with cells colored by cell types (B) Density plot showing cell type specific markers for vascular smooth muscle cell (*ACTA2*, *MYH11*, *MYL9*, and *TAGLN*) and vascular fibroblasts (*LUM*, *DCN*, *COL3A1*, and *LAMA2*). (C) Bar plot showing significant *9p21.3* tile-to-gene pair counts of *9p21.3* cis-expressing genes (*MTAP*, *DMRTA1*, and *CDKN2B-AS1*) stratified by genotype at rs1537371 in vascular fibroblasts and smooth muscle cells (D) Genomic track shows accessibility of *9p21.3* locus at rs1537371 SNP in vascular fibroblast. (E) Allelic dependent pseudobulk mRNA expression of *MTAP* in vascular fibroblast (F) Genomic track shows accessibility of *9p21.3* locus at rs1537371 SNP in vascular smooth muscle. (G) Allelic dependent pseudobulk expression of MTAP in vascular smooth muscle

## Supplementary Tables (available as spreadsheet workbook)

**Table S1.** PheWAS data from FinnGen freeze 13, shown in panel 1A, for the *9p21* tag SNP rs4977574. *9p21.3* is robustly associated with coronary artery disease and other vascular disease traits.

**Table S2.** The haplotype table, sourced from HaploReg (v4.2)^3^, presents variant information in linkage disequilibrium with the primary *9p21* variant.

**Table S3.** LD-score regression heritability analysis of vascular disease and control (left-handedness and schizophrenia) traits in vascular cells under nutrient and inflammatory stimulatory conditions.

**Table S4.** Differential H3k27ac peaks within the *9p21* locus for 4 vascular cell lines, vascular fibroblasts, smooth muscle cells, endothelial cells, and adipose pericytes. Differential peaks were calculated using DiffBind, with a genome-wide significance level of FDR=0.05.

**Table S5.** Potentially cis-regulated target genes within the *9p21* TAD, gene function, and distance from each gene TSS to the locus.

**Table S6.** CRISPRi-MAC-Seq *9p21* library screen structure. The library contains 417 guides, including locus-targeting controls, scrambled controls, TSS, enhancer, and SNP-targeting guides. This table included guide position, sequence, the target the guide was designed for, and an additional column to note enhancer overlap of SNP-associated guides.

**Table S7**: Proportion of the 417 guides in the screen targeting each control type or genomic feature. Also shown in Figure 2c.

**Table S8.** Constraint analysis of the *9p21.3* locus. Constraint values are calculated for 1kb blocks using Gnocchi, and represent a constraint z-score relative to the whole genome. Regions that overlap SNPs from the CAD GWAS haplotype are indicated; a Gnocchi z-score of ≥ 2 was used as a threshold for significant constraint.

**Table S9.** WASP ^4^ was used to examine the link between heterozygous *9p21* variants and regulatory targets. A cutoff of P< 0.05 was considered to identify significant variant effects of individual *9p21.3* haplotype variants. Subsequently, the sum of the read counts corresponding to significant effects was used to represent the effect of the *9p21.3* haplotype on individual genes.

**Table S10.** HuGE-AMP data from the Common Metabolic Diseases Knowledge Portal, using genetic evidence from rare and common genetic variation to implicate MTAP in vascular disease processes (accompanies figure S8).

**Table S11.** Rare-variant analysis of potential *9p21.3* targets indicated nominal significance of *MTAP* (p=0.0919) in disease phenotypes.

**Table S12**. International Mouse Phenotyping Consortium (IMPC) database showing phenotypes associated with *Mtap* haploinsufficiency

**Table S13**. Gene prioritization using MAGMA. This table shows MAGMA gene-based Z-scores and P-value for CAD GWAS.

**Table S14.** Schizophrenia-risk GWAS gene-trait association for *9p21.3* genes. Intended as a control trait for CAD-risk, only one *9p21.3* gene, *MIR31HG*, ranked above the 95th percentile.

**Table S15.** DEGs (generated using DESeq2) for siMTAP knockdown relative to the non-targeting control in vascular smooth muscle cells under basal conditions.

**Table S16.** Gene Ontology (GO) enrichment of the differentially expressed genes (DEG) in VSMC was performed to identify functional clusters of enriched pathways. This table presents the mapping of pathways to clusters and provides functional annotations for each cluster.

**Table S17.** Network properties for functional clusters enriched in differentially expressed genes in vascular smooth muscle cells.

**Table S18.** DEGs from comparing basal vs MTAP perturbation in vascular fibroblasts

**Table S19.** Gene Ontology (GO) enrichment of the differentially expressed genes (DEG) in fibroblast was performed to identify functional clusters of enriched pathways. This table presents the mapping of pathways to clusters and provides functional annotations for each cluster.

**Table S20.** Network properties for functional clusters enriched in differentially expressed genes in vascular fibroblasts.

**Table S21**. Total number of morphological extracted features captured by LypocyteProfiler on vascular fibroblast and smooth muscle cells

**Table S22**. Pearson correlation of LipocyteProfilerfeatures in vascular fibroblasts across individual replicates of treatment conditions.

**Table S23**. Pearson correlation of LipocyteProfilerfeatures in vascular SMCs across individual replicates of treatment conditions.

**Table S24.** LipocyteProfiler feature comparison between gene control and siMTAP treatment in vascular fibroblasts for basal and TGF-β stimulated conditions

**Table S25.** LipocyteProfiler feature comparison between gene control and siMTAP treatment in vascular fibroblasts for basal and TGF-β stimulated conditions.

**Table S26.** MAC-seq codes used for multiplex library preparation and sequencing.

**Table S27.** Statistical analysis (Wilcoxon) of pseudobulked, annotated cell types from a human dilated and hypertrophic cardiomyopathy dataset.

**Table S28.** Quantification and statistical details for image analysis QC

**Table S29.** Primers used for qPCR analysis of *9p21.3* genes.

**Table S30**. Patient demographies for sc-Multiome

**Table S31.** Output from SCENT analysis for vascular fibroblasts and smooth muscle cells

**Table S32.** Genome-wide distribution of MAGMA gene-based Z-scores and P-value for T2D GWAS

